# Human Histone Fragments Display Antibacterial Properties against *Pseudomonas aeruginosa*

**DOI:** 10.64898/2026.05.11.724237

**Authors:** Nourice Jaber, Angela Di Somma, Armando A. Rodríguez-Alfonso, Carolina Cané, Clarissa Read, Ludger Ständker, Sebastian Wiese, Angela Duilio, Jan Münch, Barbara Spellerberg

## Abstract

**Background:** Rising antimicrobial resistance rates, require new therapeutic approaches such as antimicrobial peptides (AMPs), which are part of the innate immune defense, as alternatives to antibiotics. In this study, we aim to unravel the antibacterial activity of human histone H1.2 peptide against *Pseudomonas aeruginosa* and its potential immune modulatory role.

**Methods:** We used a hemofiltrate peptide database for antimicrobial peptide prediction to identify novel human AMPs. Thirteen sequences of histone H1 were identified as putative AMPs, synthesized, and tested against bacterial ESKAPE pathogens in a radial diffusion assay. SYTOX green assay, electrophoretic mobility shift assay, and differential proteomics assays were conducted to determine the mode of action of H1.2 peptide fragment. A crystal violet assay was performed to evaluate the inhibition of biofilm formation. The cytotoxicity of the peptide was tested in LDH and Alamar assays. Finally, to visualize the contributions of H1.2 in NETs formation, scanning electron microscopy was performed.

**Results:** The H1.2 peptide inhibited the growth of *P. aeruginosa* in a dose and pH-dependent manner without cytotoxicity towards mammalian THP-1 cells. It acts on intracellular targets to inhibit the growth of *P. aeruginosa*. STRING analysis from the differential proteomics assay showed that H1.2 targets the downregulation of proteins involved in the biogenesis of outer membrane proteins, including the folding and trafficking of outer membrane proteins across the cytoplasmic membrane. Scanning electron microscopy images showed that H1.2 forms NET-like structures capable of trapping and immobilizing *P. aeruginosa*.

**Conclusion:** The characterized antimicrobial activity of H1.2 points to a role for human histone H1 fragments in innate immunity and may represent a promising approach for the development of novel antibacterial therapies.

**Graphical Summary:** 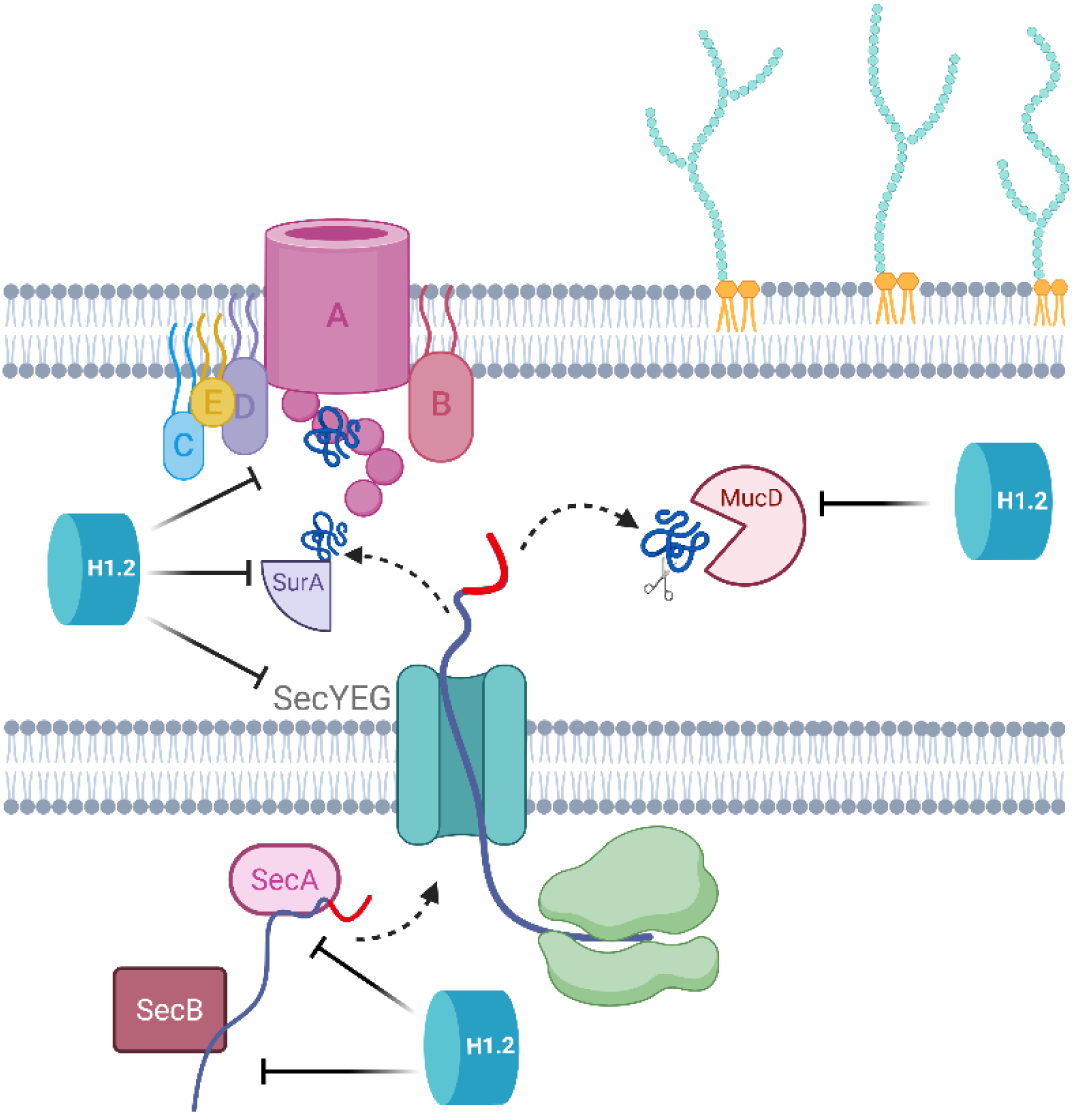

*Sec transport and BAM complex system including chaperone proteins and quality control proteases are inhibited by H1.2 in *Pseudomonas aeruginosa*.:* Outer membrane proteins (OMPs) are synthesized in the cytoplasm and transported across the inner membrane via the Sec translocase, assisted by SecA/SecB or ribosomes. In the periplasm, they are escorted by chaperones such as SurA to the BAM complex for insertion into the outer membrane. Here, we show that H1.2, an antimicrobial peptide, targets membrane biogenesis in *P. aeruginosa* through downregulating Sec translocase (SecA/SecB and SecYEG), SurA, and BAM complex. Therefore, leading to improper transfer, folding and insertion of OMPs into the outer membrane. Normally, misfolded proteins are degraded by the protease MucD to prevent toxic aggregation in the bacteria. However, with H1.2 inhibiting MucD the proteotoxic stress is exacerbated, ultimately compromising bacterial homeostasis and viability. Figure created using BioRender.com.

## Introduction

Antimicrobial resistance (AMR) represents a critical and increasing global health threat, with an estimated 1.27 million deaths directly correlated to drug-resistant infections in 2019. If current trends persist, AMR is projected to cause up to 10 million deaths annually by 2050, matching the global mortality rate of cancer (1),(2). In response to this crisis, alternative therapeutic strategies are urgently needed. Among the most promising candidates are antimicrobial peptides and proteins (AMPs), which are key components of the innate immune system and constitute effective alternatives to antibiotics. Due to their broad-spectrum bactericidal properties, these natural templates not only help prevent the onset of several infections but also enhance the efficacy of conventional antibiotics through synergistic interactions, particularly against multidrug-resistant organisms (3),(4),(5),(6). Their cationic nature and relatively small size (typically 10–50 amino acids) enable them to bind readily to negatively charged membranes, causing disruption of the membrane integrity and pore formation, eventually leading to bacterial cell lysis (3),(7),(8).

In addition to membrane-dependent antibacterial activity, some AMPs exert another direct mechanism of action by penetrating the bacterial membrane and translocating into the cytoplasm, thus targeting vital intracellular processes (9),(10). These include the inhibition of nucleic acid and cytoplasmic protein synthesis, disruption of cell wall synthesis, interference with replication, transcription, translation, or protein folding, thereby contributing to bacterial cell death through multiple complementary mechanisms (11),(12),(13).

Additionally, to their direct microbial killing, AMPs in vertebrates can indirectly clear pathogenetic infections. They play a crucial role in modulating immune responses by regulating host gene expression, acting as chemokines or stimulating chemokine production, suppressing pro-inflammatory cytokine release, promoting wound healing, and recruiting adaptive immune cells into the site of infection. Through these functions, AMPs are considered key mediators in bridging both innate and adaptive immunity together (9),(12),(14).

An interesting subfamily of AMPs is histones, small, positively charged proteins consisting of linker histone (H1) and core histones (H2A, H2B, H3, and H4), that were traditionally recognized for their role in binding to and structuring DNA within chromatin. Histones were first reported to possess antibacterial activity as early as 1942, in studies using extracts from calf thymus (15). Extranuclear histones are found in the cytoplasm, on the cell membrane, in extracellular fluid, and within neutrophil extracellular traps (NETs), where they contribute to host defense mechanisms (16),(17). NETs are web-like structures composed of decondensed chromatin and bactericidal proteins that are released from active neutrophils during infection or inflammation in a process known as NETosis. The DNA-protein fibers can trap and immobilize both Gram-positive and Gram-negative bacteria, ultimately contributing to bacterial cell death (18),(19). Histones represent the major protein component of NETs, accounting for around 70% of their total protein content, with H2A and H2B being the most abundant (20),(21). Based on this natural mechanism, synthetic NET-like structures, termed synthetic microwebs, have recently been developed to further evaluate their antimicrobial performance with varying DNA/histone compositions. Higher histone content incorporated into synthetic microwebs was shown to increase electrostatic interactions with bacterial surfaces, thereby enhancing microbial trapping and cell wall distortion (22),(23).

In line with their significant antimicrobial properties, cytoplasmic H1 and its fragments identified in human ileal mucosa help prevent microbial translocation across the intestinal epithelium and contribute to antimicrobial defense against *Salmonella Typhimurium* (17). Additionally, histones H2A and H2B in human placenta, as well as H1 in brain tissues, can neutralize lipopolysaccharide (LPS) (24),(25). In the skin, human sebocytes abundantly express histone H4, which exhibits a potent antimicrobial activity against *Staphylococcus aureus* and *Propionibacterium acnes*, two of the most common skin-associated pathogens (26). Collectively, the detection of histones with antibacterial properties in diverse tissues emphasizes their broad contribution to innate immune defense beyond the nucleus, reinforcing their role as functional antimicrobial peptides in various organ systems (27). However, despite these findings, the mechanistic details of these bactericidal activities were not dealt with and remain largely unclear.

In this study, by combining hemofiltrate peptide library screening with computational AMP prediction, we identified thirteen fragments from human histone H1 variants H1.0, H1.2, H1.3, H1.4, and H1.5 exhibiting selective antimicrobial activity against *P. aeruginosa*. Among these fragments, the H1.2 peptide displayed the highest potency in inhibiting the growth of *P. aeruginosa* without disrupting the bacterial membrane and was then selected for further characterization. The global effect of the H1.2 fragment on *P. aeruginosa* was then investigated through differential proteomic approaches, demonstrating that the peptide inhibits bacterial growth by promoting the specific downregulation of cytosolic proteins involved in the transport, membrane insertion, and assembly of Outer Membrane Proteins (OMPs). Furthermore, our findings reveal a previously unrecognized contribution of noncytotoxic H1.2 in innate immunity through the formation of web-like structures that trap bacteria, mimicking the NETs formed by neutrophils. These findings provide new insights into the elucidation of the antimicrobial properties of histone-derived peptide H1.2 against *Pseudomonas aeruginosa,* opening the way to further studies to define its effective mechanism of action at the molecular level and its role as an endogenous innate immune effector.

## Materials and Methods

### 1. Synthesis of Histone Peptide Fragments

All histone fragments were obtained from PSL Heidelberg (PSL, Heidelberg, Germany) using F-moc chemistry. All peptides were purified to greater than 95% homogeneity by reversed-phase HPLC and diluted in double-distilled water prior to use in the biological evaluation.

### 2. Bacterial Culture

Bacterial strains used in this study are listed in Table 1. For liquid culture, *Escherichia coli*, *Pseudomonas aeruginosa*, and *Acinetobacter baumannii* were grown in lysogeny broth (LB-Miller) at 37°C with shaking (160 rpm) under aerobic conditions. *Methicillin-Resistant Staphylococcus aureus*, *Klebsiella pneumoniae*, and *Enterococcus faecium* were cultured in THY medium (Todd-Hewitt Broth [Oxoid] supplemented with 0.5% yeast extract [BD, Miami, FL]) at 37°C in a 5% CO₂ atmosphere.

**Table 1.** List of bacterial strains used and their sources.

| Bacterial strain | Source |
| --- | --- |
| *Pseudomonas aeruginosa* (BSU 856) | ATCC 27853 |
| *Escherichia coli* (BSU 1286) | Pat. S491510 Koc Azra, 17.11.11 |
| Methicillin-Resistant *Staphylococcus aureus* (BSU 1348) | ATCC 43300 |
| *Klebsiella quasipneumoniae* (BSU 1353) | ATCC 700603 |
| *Acinetobacter baumannii* (BSU 1514) | ATCC 19606 |
| *Enterococcus faecium* VRE (BSU 1516) | DSM 17050 |
| *Pseudomonas aeruginosa* (BSU 1294) | Clinical isolate, Ulm collection |
| *Pseudomonas aeruginosa* (BSU 1295) | Clinical isolate, Ulm collection |
| *Pseudomonas aeruginosa* (BSU 1296) | Clinical isolate, Ulm collection |
| *Pseudomonas aeruginosa* (BSU 1297) | Clinical isolate, Ulm collection |
| *Pseudomonas aeruginosa* (BSU 1298) | Clinical isolate, Ulm collection |
| *Pseudomonas aeruginosa* (BSU 1453) | Clinical isolate, Ulm collection |
| *Pseudomonas aeruginosa* (BSU 1454) | Clinical isolate, Ulm collection |
| **Plasmids** |  |
| PAT 28 | Patrick Trieu-Cuot, France |

#### 2.1. Radial Diffusion Assay

To evaluate the antibacterial activity of histone 1 purified peptides, a radial diffusion assay (RDA) was conducted. Overnight bacterial cultures were pelleted by centrifugation at 3000 x g and washed in 10 mM sodium phosphate buffer. Following another centrifugation, the pellet was resuspended in 10 mM sodium phosphate buffer, and the optical density was determined spectrophotometrically at O.D. 600 nm. Bacterial density was adjusted to seed approximately 2 × 10^7^ bacteria into 15 ml of 1% agarose. Plates were poured and allowed to cool for 30 min at 4°C, then 2-3 mm holes were created in the agarose using sterile wide-pore pipette tips (Axygene, a Corning brand). 10 µl of each histone fragment of the desired concentration was filled into the holes. Plates were incubated at 37°C in ambient air for 3 hours, then overlaid with 10 ml tryptic soy agar. After overnight incubation at 37°C in 5% CO2, visible clear inhibition zones around the holes were measured.

To assess the specificity of the histone fragments to *P. aeruginosa*, another set of RDA was performed using the peptide fragments tested against 7 different clinical isolates of *P. aeruginosa*. To determine the effect of pH on the antimicrobial efficacy of the 13 histone 1 fragments, a pH RDA was performed. The agarose and overlay agar were adjusted to various pH levels (5, 5.5, 6, 6.5, and 7) using NaOH and HCl before performing the experiment. All conditions were tested in five independent biological replicates for positive results and three for the negative results.

#### 2.2. Bacterial Survival Assay

The antimicrobial activity of the H1.2 histone fragment in a liquid environment was evaluated using a bacterial survival assay. Overnight bacterial cultures were diluted to an optical density (OD) of 0.1 at 600 nm, and 1 mL aliquots were centrifuged at 17,000 × g for 1 minute. The resulting bacterial pellets were resuspended in 10 mM phosphate buffer. A 90 µL volume of the bacterial suspension was mixed with 10 µL of H1.2 histone fragment to achieve final concentrations ranging from 12.5 to 100 µg/mL.

Samples were incubated at 37 °C for 24 hours, and aliquots were collected at 0, 1, 2, and 24-hour time points. Collected aliquots were serially diluted in phosphate buffer and plated on blood agar. Following overnight incubation at 37 °C in 5% CO₂, colonies were enumerated, and bacterial viability was expressed as colony-forming units per milliliter (CFU/mL). The experiment was conducted in five independent biological replicates.

### 3. Assessment of Biofilm Inhibition

#### 3.1. Evaluation of Bacterial Cell Viability

Overnight bacterial cultures grown in lysogeny broth (LB-Miller) and Tryptic Soy Broth (TSB) media were adjusted to an optical density (OD) of 0.01 at 600 nm. Bacterial suspensions in either LB or TSB were then incubated with increasing concentrations of H1.2 (12.5, 25, 50, and 100 µg/mL). Aliquots of 100 µL from each condition were transferred into sterile, non-treated polystyrene 96-well plates (Corning), with each condition tested in triplicate. Wells containing untreated bacteria served as negative controls, while wells containing only media (LB or TSB) were used as blanks.

Plates were incubated for 24 hours at 37 °C with 5% CO₂ under static conditions. After incubation, wells were washed three times with 200 µL of 1× phosphate-buffered saline (PBS). Subsequently, 100 µL of Alamar Blue Cell Viability Reagent (Thermo Fisher Scientific), diluted in the respective medium, was added to each well. Absorbance was immediately measured at 572 nm, with 600 nm as the reference wavelength, using a Tecan Infinite M200 microplate reader. Measurements were repeated at hourly intervals for a total of 3 hours, with the plate maintained at 37 °C throughout the assay period.

#### 3.2. Biofilm Inhibition Assay

Following assessment of bacterial cell viability under H1.2 treatment, wells in the 96-well plate were washed three times with 200 µL of 1× phosphate-buffered saline (PBS) to remove non-adherent cells. The plate was then air-dried for 30 minutes at room temperature. Wells were subsequently stained with 0.1% (w/v) crystal violet solution for 30 minutes, followed by two washes with 280 µL of 1× PBS to remove excess stain. Stained biofilms were then solubilized in 100% ethanol by incubating the plate at 4 °C for 10 minutes with gentle shaking at 30 rpm.

A 150 µL aliquot of the solubilized crystal violet-ethanol solution was transferred to a fresh 96-well plate, and absorbance was measured at 562 nm using a Tecan Infinite M200 microplate reader. Biofilm biomass was quantified by comparing the absorbance of treated samples to that of the untreated control, which was set at 100%. All conditions were tested in three independent biological replicates.

### 4. Cytotoxicity Assessment

#### 4.1. Cell Culture

THP-1 human monocytic cells (ATCC) were cultured in T75 cell culture flasks in 20 mL of complete RPMI-1640 medium supplemented with 10% fetal bovine serum (FBS) and 1% penicillin-streptomycin. Cells were maintained at 37 °C in a humidified incubator with 5% CO₂. Cultures were routinely passaged and maintained at a density of approximately 1 × 10⁶ cells/mL

#### 4.2. Treatment of THP-1 cells

For cytotoxicity testing, THP-1 cells were seeded into 96-well plates and treated with increasing concentrations of H1 histone peptide fragments (12.5, 25, 50, and 100 µg/mL) for 24 hours at 37 °C in 5% CO₂. The total volume per well was adjusted to 100 µL. Cells treated with 1% Triton X-100 served as a negative (cytotoxic) control, while untreated cells in culture medium served as a positive (viability) control. Wells containing medium and peptides without cells were included as background controls for normalization. Each condition was tested in triplicate across five independent experiments.

#### 4.3. Viability Assay

Cell viability was assessed using the Alamar Blue Cell Viability Reagent (Thermo Fisher Scientific). After 24-hour peptide treatment, cells were centrifuged at 12,000 × g for 10 minutes, and 25 µL of the supernatant was collected for lactate dehydrogenase (LDH) analysis. The remaining pelleted cells were resuspended in culture medium containing 25 µL of Alamar Blue reagent. Fluorescence intensity was immediately measured using a Tecan Infinite M200 microplate reader at 572 nm with a reference wavelength of 600 nm. Measurements were repeated hourly for three hours, with the plate maintained at 37 °C throughout the assay.

#### 4.4. LDH Release

Cytotoxicity was further evaluated by quantifying extracellular LDH release from treated THP-1 cells. In a separate 96-well plate, 75 µL of fresh culture medium was mixed with 25 µL of previously collected supernatant from each condition. Subsequently, 100 µL of the prepared LDH reaction mixture (TaKaRa) was added to each well. Plates were incubated in the dark at room temperature for 30 minutes. Absorbance was measured at 492 nm with a reference wavelength of 600 nm using a Tecan Infinite M200 microplate reader (Tecan group).

### 5. Mode of Action

#### 5.1. SYTOX Green Membrane Permeabilization Assay

To evaluate the effect of the H1.2 histone peptide on the membrane integrity of *Pseudomonas aeruginosa*, a SYTOX Green membrane permeabilization assay was performed. An overnight culture of *P. aeruginosa* was reinoculated into fresh medium and grown to an optical density (OD) of 0.1 at 600 nm. Bacterial cells were harvested by centrifugation at 10,000 × g for 2 minutes and the pellets were resuspended in 0.9% NaCl containing 0.5 µM SYTOX Green dye (Invitrogen).

A total of 90 µL of the SYTOX-stained bacterial suspension was transferred into each well of a black 96-well plate, followed by the immediate addition of 10 µL of H1.2 peptide at a final concentration of 100 µg/ml. Fluorescence was measured immediately using a Tecan Infinite M200 microplate reader with an excitation wavelength of 488 nm and corresponding emission settings. Bacterial cells treated with 70% ethanol served as a positive control for membrane permeabilization, while cells treated with distilled water or left untreated served as negative controls. All conditions were tested in triplicate across five independent experiments.

#### 5.2. Electrophoretic Mobility Shift Assay

Plasmid PAT28 was isolated using the QIAprep Spin Miniprep Kit (Qiagen, 250 preps) and digested with the restriction enzyme BamHI (New England Biolabs) according to the manufacturer’s protocol. The digested plasmid DNA was subsequently purified using the QIAquick PCR Purification Kit (Qiagen).

For the electrophoretic mobility shift assay (EMSA), 100 ng of purified plasmid DNA (*E. coli* PAT) 28 was incubated with increasing concentrations of the H1.2 histone peptide (0, 2, 5, 10, 50, and 100 µg/mL) in a total volume of 30 µL Tris-EDTA (TE) buffer. The mixtures were incubated at room temperature for 30 minutes to allow for DNA–peptide complex formation. The native loading buffer was then added to each reaction (28).

A 20 µL aliquot of each sample was loaded onto a 1.5% agarose gel, and electrophoresis was performed under non-denaturing conditions at 160 V for 1 hour. DNA migration was visualized by staining with ethidium bromide and imaged under UV illumination. A reduction in DNA mobility through the SDS gel indicates complex binding between the H1.2 peptide and the plasmid DNA. This experiment was tested in three independent replicates.

#### 5.3. Proteomics Assay

##### 5.3.1. Bacterial Preparation

An overnight culture of the bacterial strain *P. aeruginosa* (BSU 856) was reinoculated into 100 mL of fresh lysogeny broth (LB-Miller) and incubated overnight at 37 °C with shaking at 180 rpm. The following day, the culture was adjusted to an optical density (OD₆₀₀) of 0.5 in 25 mL of LB medium and treated with the H1.2 histone fragment at a final concentration of 25 µg/mL. The suspension was incubated for 3 hours at 37 °C under the same shaking conditions.

After incubation, the bacterial suspension was centrifuged at 5,000 rpm for 20 minutes at 4 °C. The supernatant was carefully discarded, and the resulting bacterial pellet was stored at - 80 °C until further analysis.

##### 5.3.2. Differential Proteomics Assay

Briefly, bacterial pellets were resuspended in phosphate buffer saline (PBS), 5% sodium dodecyl sulfate (SDS), and 1 mg/mL lysozyme, then lysed by sonication. Samples were then centrifuged at 4 °C for 30 min at 13000 rpm to collect the unlysed cells and cell debris, while the recovered supernatant was quantified using the Pierce™ BCA Protein Assay Kit purchased from Thermo Scientific (Rockford-USA). 50 µg of each sample were digested with trypsin onto S-trapTM micro spin column, according to the manufacturer’s instructions (Protifi, Huntington, NY). The obtained peptide mixtures were analyzed by LC-MS/MS using an LTQ Orbitrap XL coupled to a nanoLC system (ThermoFisher Scientific, Waltham, MA). All peptide mixtures were fractionated onto a C18 capillary reverse-phase column (200 mm length, 75 μm ID, 5 μm biosphere), using a non-linear 5%–50% gradient for eluent B (0.2% formic acid in 95% acetonitrile) in A (0.2% formic acid and 2% acetonitrile in MilliQ water) over 260 min. MS/MS analyses were performed in Data-Dependent Acquisition (DDA) mode by fragmenting the 10 most intense ions in Collision-induced dissociation (CID) modality. All samples were run in duplicates. The obtained data were analyzed with MaxQuant (v.1.5.2.8) using UniProt *P. aeruginosa* as a database for Andromeda search (29). Proteins were identified based on two or more peptides, at least of which was unique. Methionine oxidation and pyroglutamate formation at the N-terminal glutamine were considered as variable modifications. The accuracy for the first search was set at 10 ppm, then lowered to 5 ppm in the main search. A False Discovery Rate (FDR) of 0.01 was used, with a reverse database as decoy; retention time alignment and second peptide search functions were allowed. Fold changes (FCs) were calculated according to Label Free Quantification (LFQ) values (30).

Also, protein-protein interaction (PPI) networks were analyzed using the STRING program version 12.0 (string-db.org), which predicts PPI by employing a combination of prediction approaches and experimental data (textmining, experiments, databases, co-expression, neighborhood, gene fusion, co-occurrence).

#### 5.4. Reverse Transcription Polymerase Chain Reaction

To further follow up with the downregulation of outer membrane proteins at the transcriptional level, quantitative analysis of RNA expression was performed using reverse transcription PCR (RT-PCR). Overnight bacterial cultures were diluted and grown to an optical density (OD₆₀₀) of 0.5, followed by the treatment with 25 µg/mL of the H1.2 peptide for 1 hour at 37 °C. Untreated bacterial cultures served as negative controls.

Total RNA was extracted using the RNeasy Mini Kit (Qiagen) according to the manufacturer’s instructions. Residual genomic DNA contamination was eliminated by DNase I digestion (Roche, Cat. No. 4716728001). RNA concentration and purity were assessed using the Qubit RNA HS Assay Kit (Thermo Fisher Scientific), and samples were diluted with RNase-free water to final concentrations of 10, 5, 2.5, 1.25, and 0.625 ng/µL.

One-Step RT-PCR reactions (Qiagen) were prepared using gene-specific primers targeting *secY*, *secA*, and the housekeeping gene *rpsL30* (primers sourced from Biomers.net). Reactions were performed using each RNA concentration as a template. The thermal cycling was performed on a PCR cycler with a protocol optimized for the OneStep RT-PCR kit.

Genomic DNA from *P. aeruginosa* served as a positive control, and nuclease-free water was used as a no-template negative control. To confirm complete DNase treatment, reactions were performed in parallel with and without reverse transcriptase.

Following amplification, ethidium bromide-containing loading buffer was added to the PCR products, which were subsequently resolved by agarose gel electrophoresis and visualized under UV illumination.

### 6. Preparation of Artificial Microwebs and Bacterial Entrapment for Scanning Electron Microscopy

To determine the efficacy of H1.2 in forming artificial microwebs and trapping bacteria, H1.2 peptide was mixed with phage DNA (Sigma, D3779) and *P. aeruginosa*, and then visualized by scanning electron microscopy (SEM).

To prepare the microwebs for SEM, a poly-L-lysine-coated silicon wafer was placed in a single well of an 8-well chamber slide (SARSTEDT). In a total volume of 20 µl, HBSS Hank’s Balanced Salt Solution (Sigma, 55037C-1000 ml) was added onto the silicon wafer, followed by the addition of methylated lambda phage DNA (Sigma, D3779) and the peptide at a final concentration of 166 and 866 µg/ml, respectively. As a positive control, methylated lambda phage DNA was added to Calf thymus histones (Sigma, H9250) at a final concentration of 100 and 500 µg/ml, respectively. DNA and Calf thymus histone concentrations were measured with Nanodrop (Thermo Scientific) before each preparation and diluted accordingly with HBSS (22). All samples inside the chamber wells were then sonicated in an ultrasonic bath (GEN-PROBE) for 10 seconds and incubated for 2 hours at 37 °C with agitation at 70 rpm to allow microweb formation.

For bacterial entrapment, an overnight bacterial culture of *P. aeruginosa* was reinoculated into fresh medium and grown to an O.D. of 0.1 at 600 nm. After microwebs’ sonication, bacteria were added at a final O.D. of 0.001 in a total volume of 20 µl of microwebs, and then incubated for 2 hours at 37 °C with agitation at 70 rpm.

Following incubation, the formed microwebs on the poly-L-lysine-coated silicon wafers (∼1 cm x 0.4 cm) were fixed by covering the wafers with the respective preparation with fixation solution (2.5 % glutaraldehyde, 1 % sucrose in 0.2 M phosphate buffer, pH 7.3) and kept overnight at 4 °C. Further preparation was performed as described previously (31). In detail, samples on silicon wafers were post-fixed with 2% osmiumtetroxide in PBS for 20 min, washed in PBS, gradually dehydrated with increasing propanol concentrations, critical point dried, and sputter coated with 2 nm platinum (CCU-010, Safematic) before they were imaged using a field emission scanning electron microscope (Hitachi S-5200, Tokyo, Japan) at an acceleration voltage of 5 kV.

### 7. Statistical Analysis

Statistical analysis was performed by using GraphPad Prism (Version 10.5.0). Data are presented as mean plus or minus standard deviation (SD) from at least three to five independent biological replicates. Comparisons between two groups were conducted using an unpaired two-tailed multiple Mann-Whitney U test. *p*-values < 0.05 were considered statistically significant.

## Results

### Identification of H1 peptide fragments and their antibacterial activity against ESKAPE pathogens

The novel putative antimicrobial peptides derived from human histone H1 were identified from a hemofiltrate peptide database library (Rodriguez et al. 2026, manuscript in preparation) comprising more than 13000 peptide sequences. Using CAMPR3, iAMPpred, and AMP scanner vr2, in silico predictions were carried out to predict the antimicrobial activity (32),(33),(34). A list was constructed in decreasing order of average AMP score. The top-scoring peptides were composed of 13 Histone H1 fragments belonging to the following histone H1 variants: H1.0, H1.2, H1.3, H1.4, and H1.5 (Table 2). The selected peptides were synthesized, and their antibacterial activity was assessed against ESKAPE pathogens using radial diffusion assays (RDA).

**Table 2.**
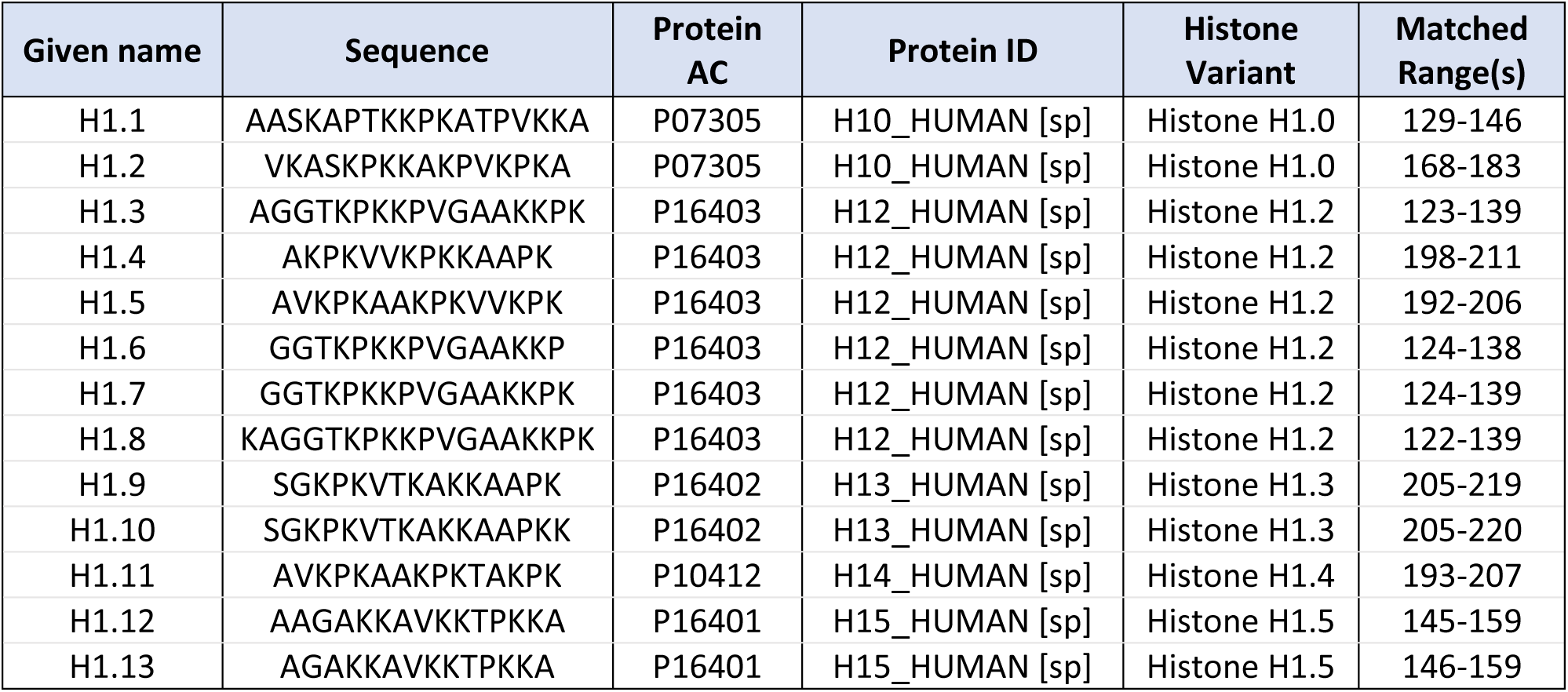
List of unmodified H1 peptide fragments. This table shows the identified H1 peptide fragments, their sequences, protein accession number (AC) in Uniport and their matched ranges within the full-length histone variants.

Strikingly, 12 out of 13 H1-derived fragments exhibited pronounced antibacterial activity against *Pseudomonas aeruginosa*, whereas no inhibitory effects were observed against *Escherichia coli*, vancomycin-resistant enterococci (VRE), *Klebsiella quasipneumoniae*, *Acinetobacter baumannii*, or methicillin-resistant *Staphylococcus aureus* (MRSA) (Figure 1A). This indicates a highly selective antibacterial profile.

**Figure 1.**
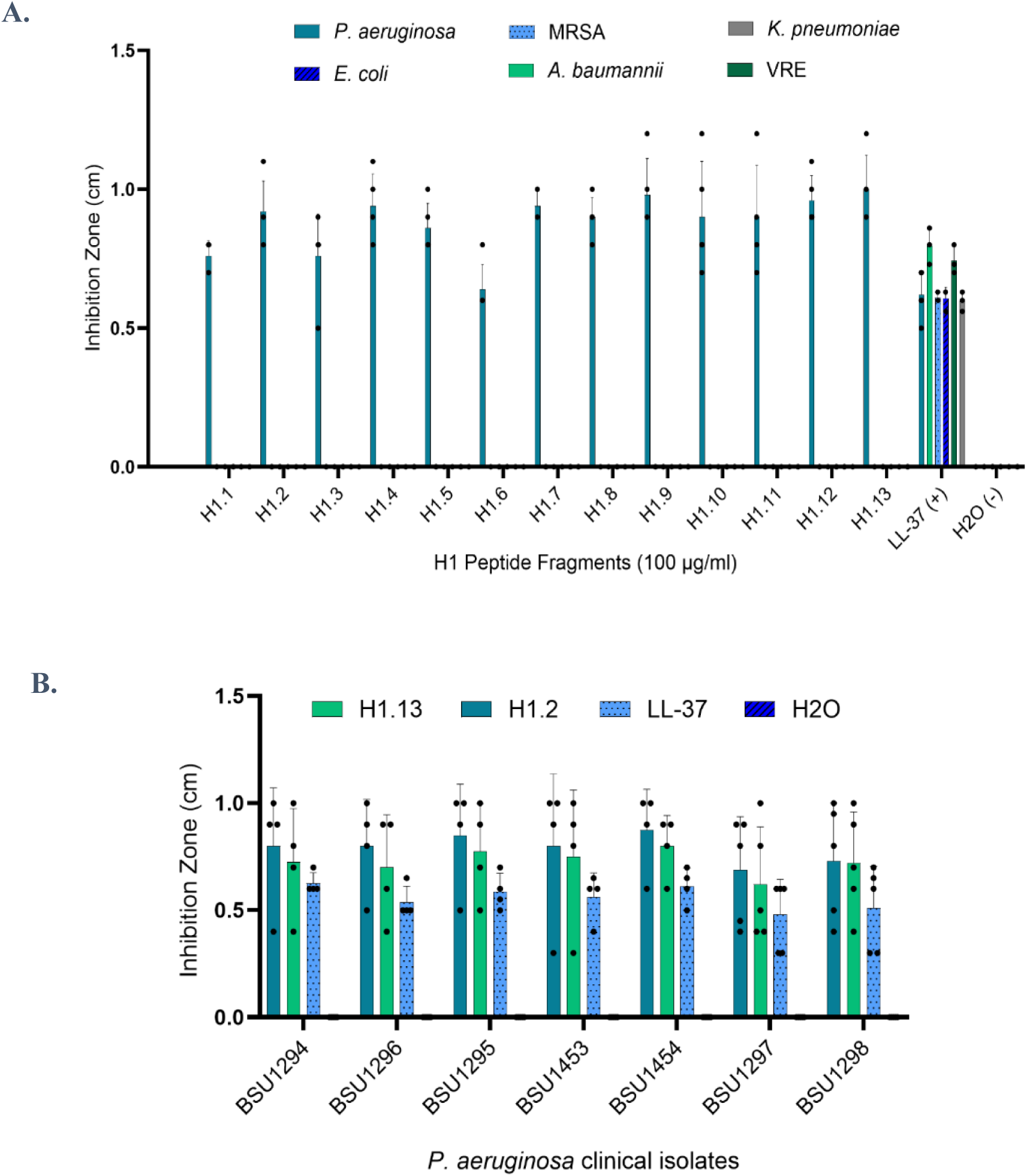
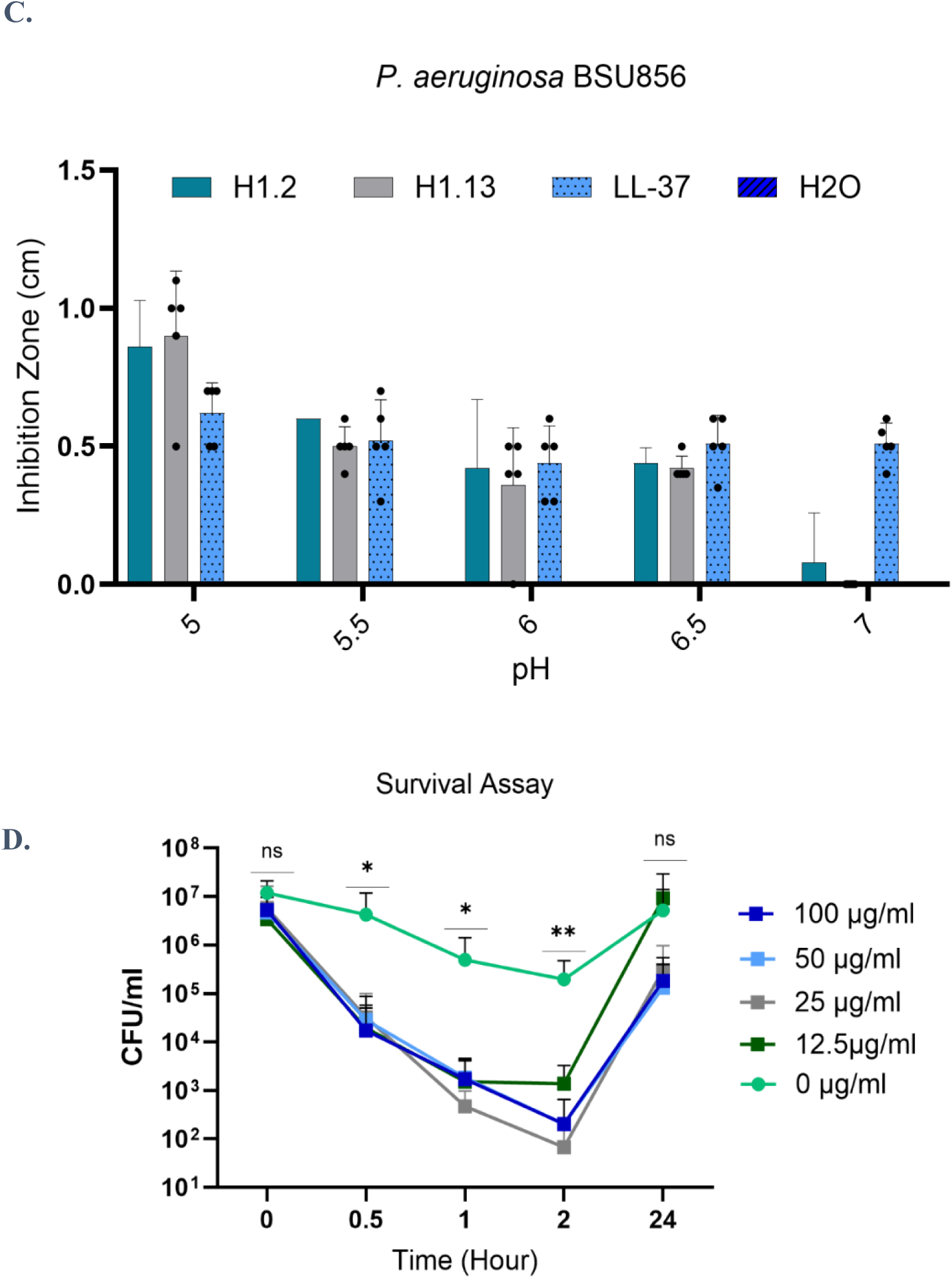
Histone peptides inhibit the growth of *P. aeruginosa*. **(A)** RDA assay shows that the 13 fragments of histone 1 peptides are specifically active in inhibiting the growth of *P. aeruginosa,* where clear inhibition zones were noticed. **(B)** The thirteen fragments of H1 peptide are active against several *P. aeruginosa* clinical isolates. **(C)** The antibacterial activity of H1.2 peptide fragment was tested in different pH media in RDA. **(D)** The effect of H1.2 peptide fragment on *P. aeruginosa* was quantified in a survival assay. In a liquid culture H1.2 reduced the bacterial survival of by 5-log_10_ after 2 hours of treatment. Values are expressed in CFU/ml.

To further investigate this specificity, the most active peptides (H1.2 and H1.13) were tested against multiple clinical isolates of *P. aeruginosa*. All tested strains were susceptible at 100 µg/ml (Figure 1B), demonstrating consistent activity across genetically diverse isolates and underscoring species-specific efficacy.

Given that infection sites often exhibit acidic microenvironments, we next evaluated peptide activity under different pH conditions. H1.2, representative of the H1 fragments, displayed maximal antibacterial activity at acidic pH, while activity was markedly reduced or absent at neutral pH (Figure 1C and S1). This suggests that environmental conditions critically modulate peptide function.

To validate these findings in a dynamic system, antibacterial activity was further assessed in liquid culture. H1.2 induced a rapid and potent bactericidal effect, reducing *P. aeruginosa* viability by approximately 3 log₁₀ within 30 minutes and up to 5 log₁₀ after 2 hours in a dose-dependent manner. Notably, partial bacterial regrowth was observed after 24 hours (Figure 1D), indicating that the effect is potent but not fully sterilizing under the tested conditions.

To enhance peptide stability in liquid environments, we evaluated the antibacterial activity of H1.2 peptide in its D-amino acid version in another RDA assay. However, this modification did not improve the activity compared to the original peptide (Figure S2). Given the comparable activity profiles, subsequent investigations of the mode of action were performed using the original H1.2 peptide fragment.

These findings identify H1 peptides as selective and pH-dependent antibacterial effectors with rapid activity against *P. aeruginosa*.

### Analyzing the influence of structural modifications on the antibacterial activity of H1 peptides

The amino acid sequence of AMPs is essential for their fundamental structure and overall antibacterial activity (35). Therefore, to further optimize the activity of H1 peptide fragments, single-point modifications were proposed for these 13 Histone fragments using CAMPR3 (rational design tool), and then evaluated with iAMPpred, AMP scanner VR2, and AI4AMP [55]. The list of modified peptides is represented in Table 3. For instance, 13-10-I is a modification of H1.13, in which Threonine (Thr) was substituted with Isoleucine (Ile) at position 10. The antibacterial activity of the modified H1 peptides was tested in a radial diffusion assay against *P. aeruginosa* in a concentration-dependent manner. Unexpectedly, the antibacterial activity of the modified fractions was reduced compared to the original H1.2 and H1.13 histone fragments (Figure 2A). To visualize this structural modification and to better relate it to antibacterial activity, a 3D structural illustration was created using the AlphaFold Protein Structure Database. H1.0 and H1.5 are the full-length variants of fragments H1.2 and H1.13, respectively. The full-length histones contain α-helical, β-sheet, and random-coil structures (Figure 2B (1) and (3)). Highlighted in green are the 3D structures of H1.2 and H1.13 peptides in Figures 2B (1) and 2B (3), respectively; these peptides are part of the random coil structures. The substitution of Thr with Ile in H1.13 at position 10 (13-10-I) didn’t change its secondary structure according to AlphaFold analysis (Figure 2B (4)). However, the molecular structure illustration highlights the difference between 13-10-I (modified fragment) and H1.13 (original fragment). Clearly, in H1.13, four hydrogen bonds are noticed (Ala 5 - Val 7), (Lys 6 - Lys 8), (Val 7 - Lys 9), and (Thr 10 - Lys 8) while in the prediction for 13-10-I two hydrogen bonds are missing upon the Ile substitution, leaving behind (Lys 5 - Val 7) and (Lys 8 - Ile 10). This suggests that the Thr substitution influenced the hydrogen bonding of the modified peptide, reducing its antibacterial activity.

**Figure 2.**
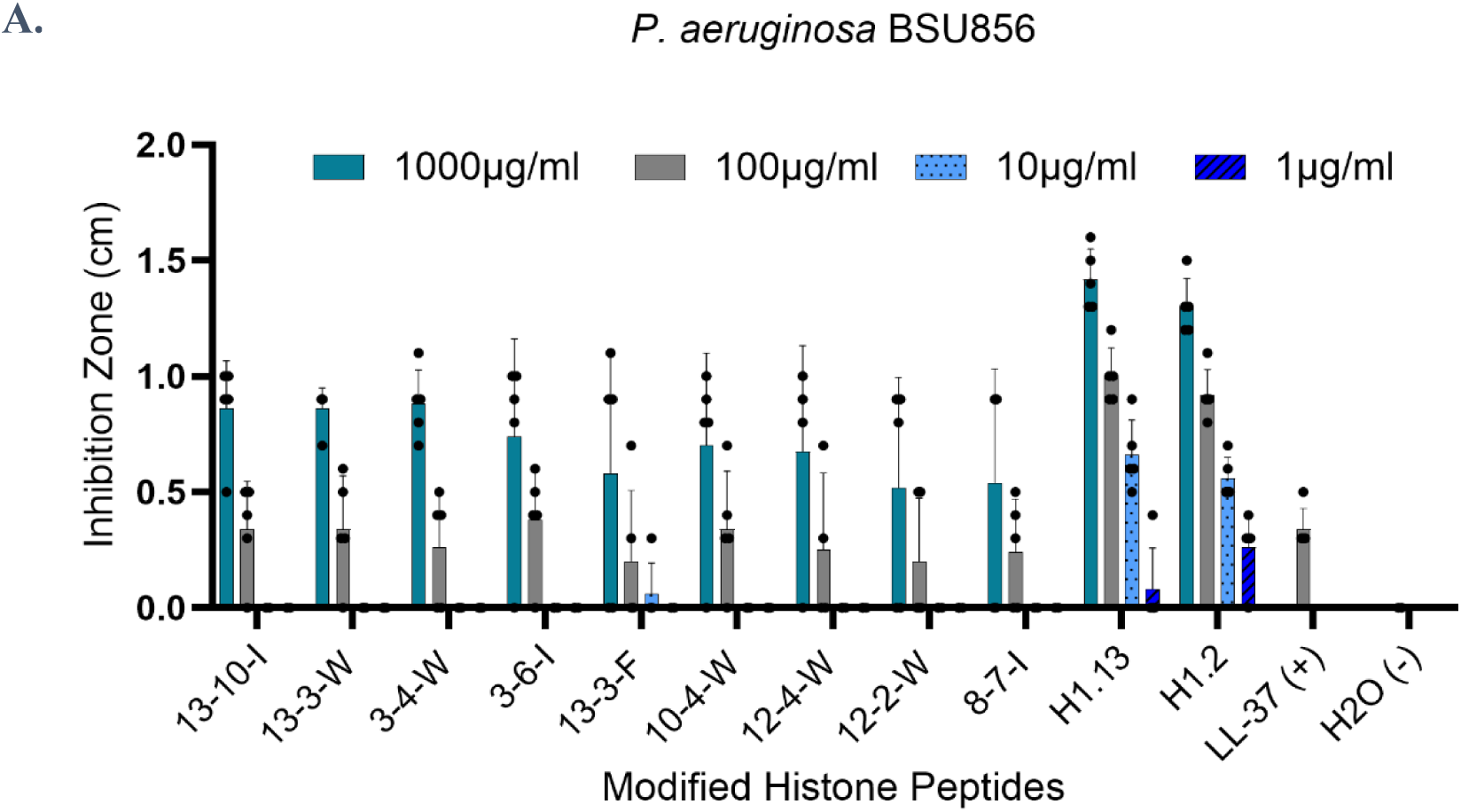
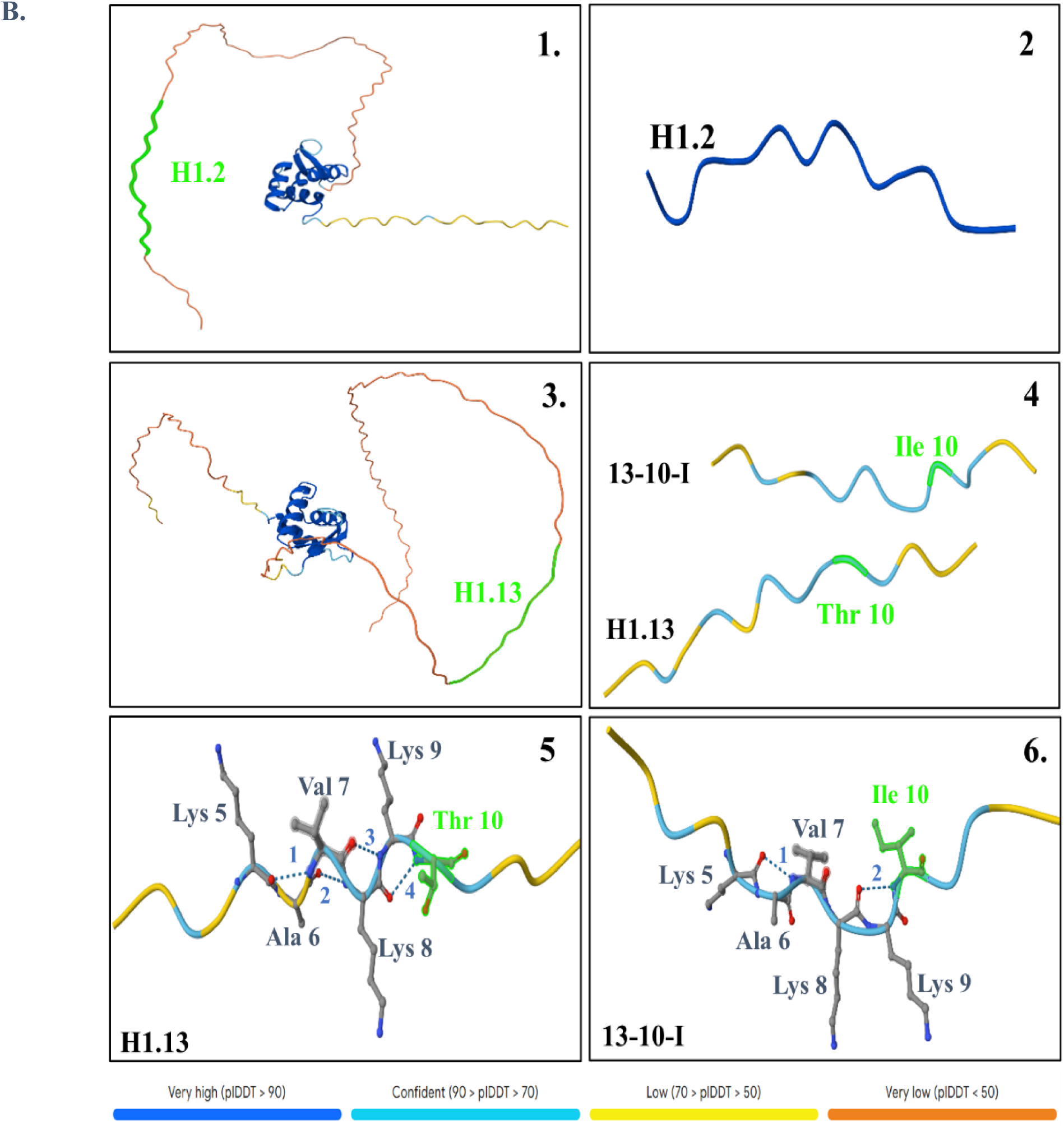
**(A)** Modified fragments of H1 peptide are less active in inhibiting the growth of *P. aeruginosa*. **(B)** 3D structure of H1.0 histone 1 variant and highlighted in pink is H1.2 peptide fragment (1) and (2). 3D structure of H1.5 histone 1 variant and highlighted in pink is H1.13 (3). 3D structure of H1.13 and its modified fragment 13-10-L that has a substitution of amino acid Thr by Ile on position 10 (4), (5) and (6). Different hydrogen bonds identified in H1.13 (5) numbered: **1:** Lys 5-Val 7, **2:** Ala 6-Lys 8, **3:** Lys 9-Val 7, **4:** Thr 10-Lys 8. Hydrogen bonds identified in 13-10-I (6) were less **1:** Val 7-Lys 5, **2**: Lys 8-Ile 10. Structures were formed with AlphaFold Server, showing the confidence score (plDDT) for each amino acid residue in the predicted structure of the residue.

**Table 3.**
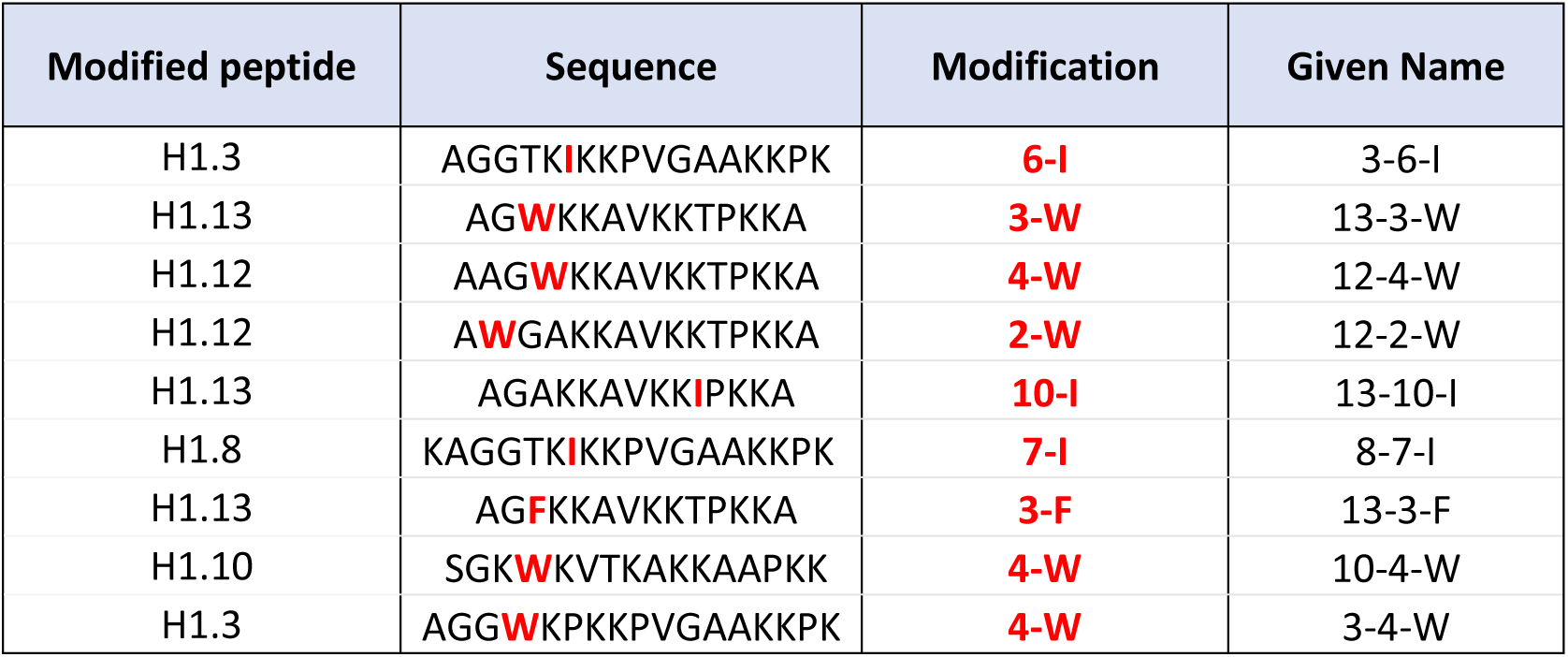
List of modified H1 peptide fragments. This table shows the list of predicted modified peptide fragments, the applied modification and their sequences.

### Assessment of H1.2 Fragment Activity Against *P. aeruginosa* Biofilm

*P. aeruginosa* is a well-established biofilm-forming pathogen, particularly in cystic fibrosis patients. To evaluate the potential anti-biofilm activity of the H1.2 histone fragment, *P. aeruginosa* cultures were treated with increasing concentrations of H1.2 (12.5, 25, 50, and 100 µg/ml) in a 96-well plate and incubated overnight at 37°C in LB or TSB media to allow biofilm formation. Bacterial cell viability within the biofilm was assessed using the Alamar Blue assay. Viability increased over time in both treated and untreated samples, and even at the highest H1.2 concentration, no reduction in *P. aeruginosa* viability was observed in either growth medium (Figures S3 A and B). In addition, a crystal violet assay was performed to quantify biofilm. The results showed no significant differences in biofilm biomass between treated and untreated samples, even at 100 µg/ml H1.2 (Figure S3 C). These findings indicate that the H1.2 histone fragment does not inhibit biofilm formation by *P. aeruginosa*.

### Cytotoxicity assessment of H1 peptides in THP-1 cells

To evaluate the potential cytotoxic effects of H1 peptides on human cells, THP-1 monocytic cells were incubated with increasing concentrations of H1.2 (12.5–100 µg/mL) for 24 hours. Cytotoxicity was assessed by measuring lactate dehydrogenase (LDH) release, while metabolic activity was evaluated using the Alamar Blue assay. H1.2-treated cells exhibited LDH levels comparable to untreated controls, indicating no detectable membrane damage, whereas Triton X-100 treatment (positive control) resulted in a marked increase in LDH release (Figure 3A). Consistently, Alamar Blue assays showed sustained metabolic activity in H1.2-treated cells, as reflected by the efficient reduction of resazurin to resorufin, comparable to control cells (Figure 3B). Together, these results demonstrate that H1.2 does not induce cytotoxic effects in THP-1 cells under the tested conditions, supporting its selective antibacterial activity.

**Figure 3.**
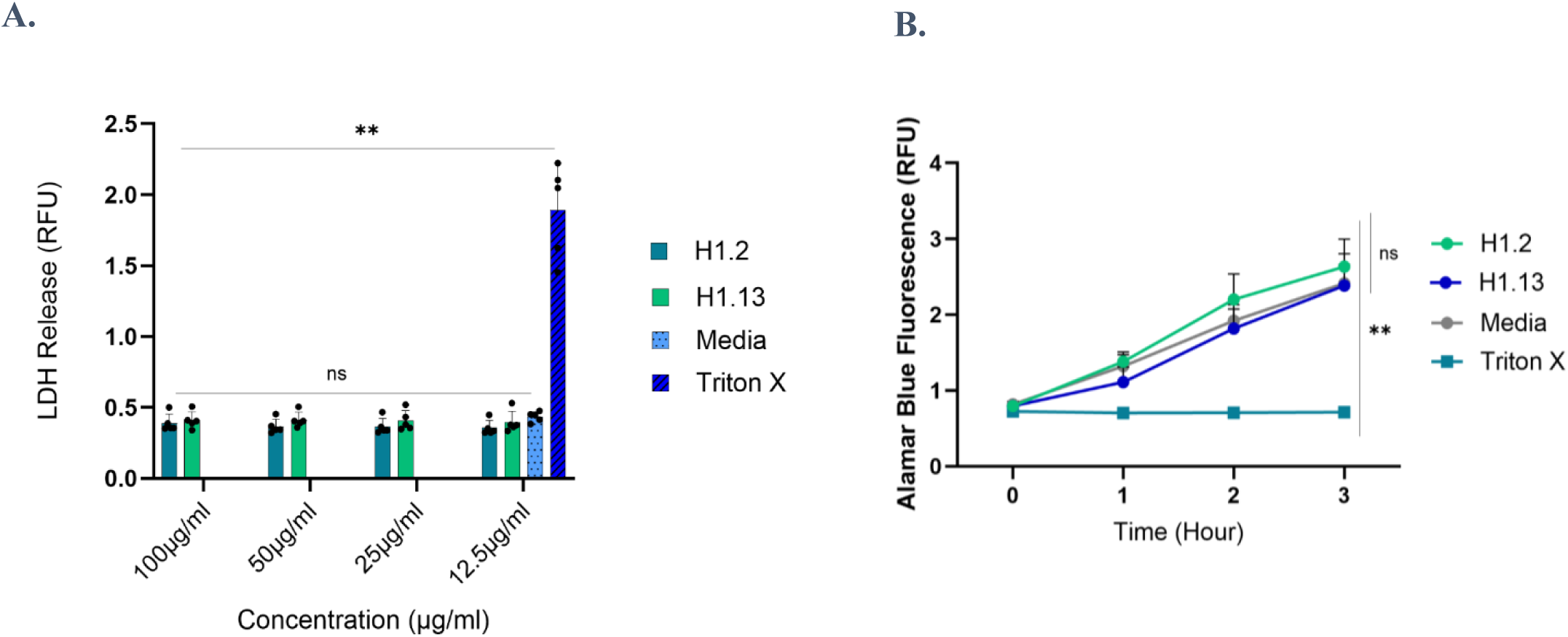
Histone peptides are not cytotoxic to mammalian THP-1 cells. **(A)** LDH assay shows that supernatant of THP-1 cells treated 24 hours with H1 peptide fragments does not induce the release of LDH in comparison to Triton X treated cells. **(B)** The Alamar Blue assay indicates that H1 peptide fragments do not interfere with the metabolic activity of THP-1 cells. Resazurin is reduced to fluorescent resorufin in both control and peptide-treated cells.

### Evaluation of membrane integrity following H1.2 treatment

Antimicrobial peptides (AMPs) exert bactericidal effects by causing bacterial membrane damage; therefore, we investigated whether H1 fragments also act through membrane disruption mechanisms. To assess membrane integrity, a SYTOX Green assay was performed. SYTOX is a green fluorescent dye that is activated when it binds to DNA. *P. aeruginosa* bacterial cells were treated with different concentrations of H1.2. Bacterial cells treated with the H1.2 peptide fragment emitted fluorescence comparable to that of the media-treated bacteria (Figure 4A), which served as a negative control, suggesting that H1.2 peptide does not target the bacterial membrane to inhibit the growth of *P. aeruginosa*.

**Figure 4.**
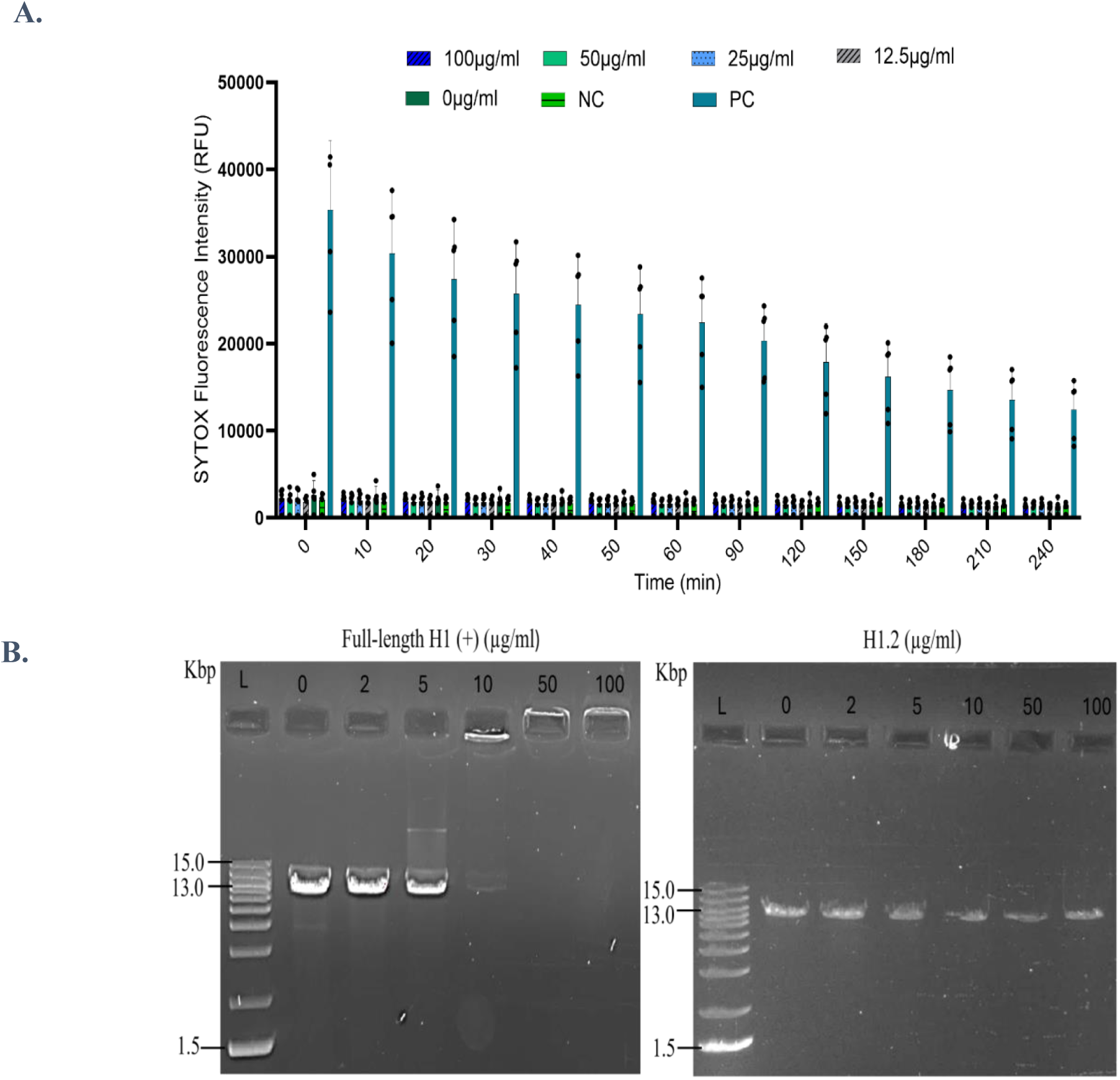
H1.2 Bacterial membrane and bacterial DNA are not targets for H1.2 histone peptide. **(A)** The fluorescence intensity of SYTOX detected shows that the incubation of different concentrations of H1.2 with *P. aeruginosa* for 3 hours does not induce any membrane disruption compared to the ethanol-treated bacteria (positive control). **(B)** DNA binding assay showing that H1.2 does not bind to bacterial DNA, while the full-length H1 (+) binds to bacterial DNA starting from 10µg/ml. (L): Ladder, Invitrogen 1Kb Plus DNA Ladder Cat.No. 10787018.

### Investigation of H1.2 Interaction with Bacterial DNA

The H1.2 fragment originates from the H1.0 histone protein, known for its DNA-binding properties. This prompted the hypothesis that the peptide may exert antimicrobial activity through direct interaction with bacterial DNA. To assess whether H1 fragments can bind to genomic DNA, thereby inhibiting *P. aeruginosa* growth, an electrophoretic mobility shift assay (EMSA) was conducted. *E. coli* plasmid DNA was incubated with varying concentrations of H1.2 fragments for 30 minutes at room temperature. The formation of peptide-DNA complexes was evaluated by monitoring the retardation of DNA migration on agarose gel electrophoresis. As expected, the full-length histone H1 protein (positive control) demonstrated substantial retardation of DNA migration, confirming its DNA-binding capacity. However, the H1.2 fragment showed no detectable effect on DNA migration patterns compared to the untreated control (Figure 4B). These findings indicate that the antimicrobial effect of the H1.2 fragment is independent of direct DNA binding.

### H1.2 induces downregulation of proteins involved in outer membrane biogenesis

The molecular mechanism underlying the antibacterial activity of H1.2 was investigated by differential proteomic approaches of *P. aeruginosa* following peptide treatment, according to the label-free quantitative procedure. Bacterial cells were exposed to H1.2 (25 µg/mL) for 3 hours, and global protein expression changes were assessed by LC–MS/MS on an Orbitrap instrument in DDA mode. Protein lysate from untreated *P. aeruginosa* cells was used as a control.

Comparative analysis revealed extensive proteomic remodeling, with 463 proteins differentially expressed, including 51 upregulated and 412 downregulated proteins. Detailed information for each identified protein, including the protein name, UniProt code, gene name, and the corresponding Fold Change (FC), is provided in Supplementary Tables 2 and 3.

A bioinformatic analysis using the STRING software was performed, and the results are shown in Figure 5 and Table 4. Among the downregulated proteins, key components of the Sec translocon system (SecA, SecD, SecF, SecY, YidC, and YajC), which mediates protein translocation across the inner membrane, were markedly reduced. In addition, periplasmic chaperones (SurA and Skp), required for proper folding and trafficking of outer membrane proteins (OMPs), as well as core components of the β-barrel assembly machinery (BamA, BamB, and Opr86), responsible for OMP insertion into the outer membrane, were also downregulated. These results indicate a coordinated downregulation of proteins involved in OMP transport and assembly, suggesting that peptide H1.2 impairs the Sec pathway and disrupts outer membrane protein biogenesis and homeostasis in *P. aeruginosa*.

**Figure 5.**
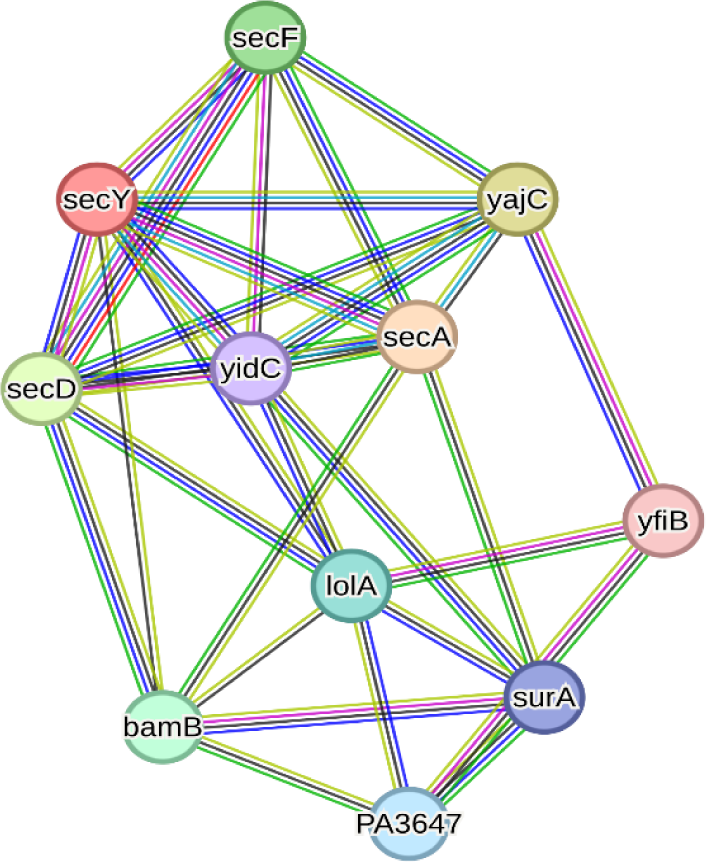
STRING network analysis of downregulated proteins following H1.2 treatment in *P. aeruginosa.* Protein-protein interaction (PPI) network scheme representing the functional clusters of putative H1.2 interactors that were significantly downregulated in the proteomic analysis. The network illustrates a high degree of connectivity among 13 nodes involved in essential pathways, specifically highlighting the integration between the Sec translocon, the BAM complex, and periplasmic chaperones.

**Table 4.**
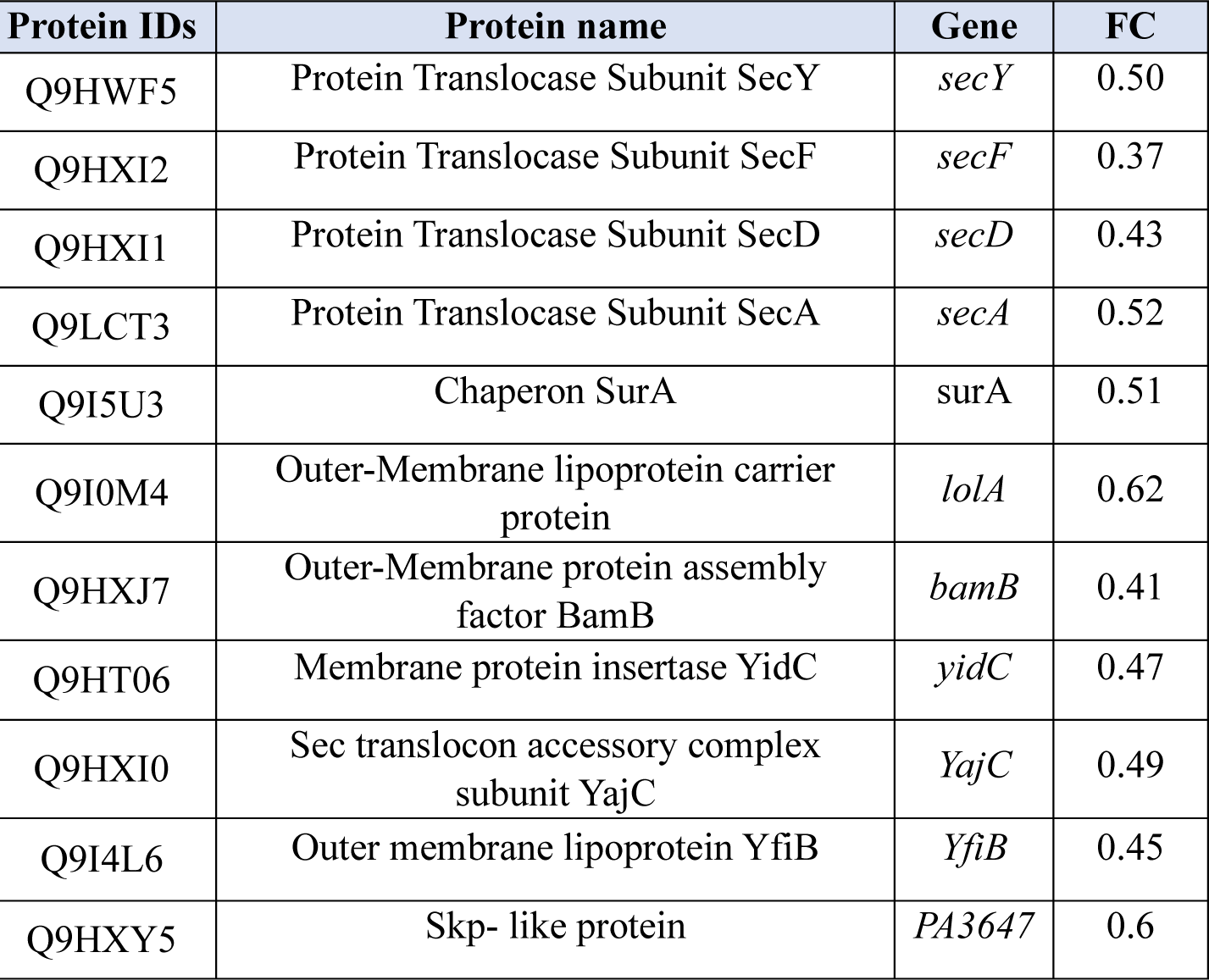
Indicating Protein IDs, Protein name, Gene, and Fold change (FC).

### H1.2 downregulates outer membrane proteins beyond transcription

To determine whether this suppression occurs at the transcriptional level, RT-PCR analysis was performed for representative genes of the Sec pathway. Despite the observed reduction at the protein level, transcripts of *SecA* and *SecY* remained detectable and comparable between treated and untreated samples across RNA dilutions (Figure 6). These findings indicate that H1.2-mediated downregulation of outer membrane biogenesis proteins occurs predominantly at a post-transcriptional level.

**Figure 6.**
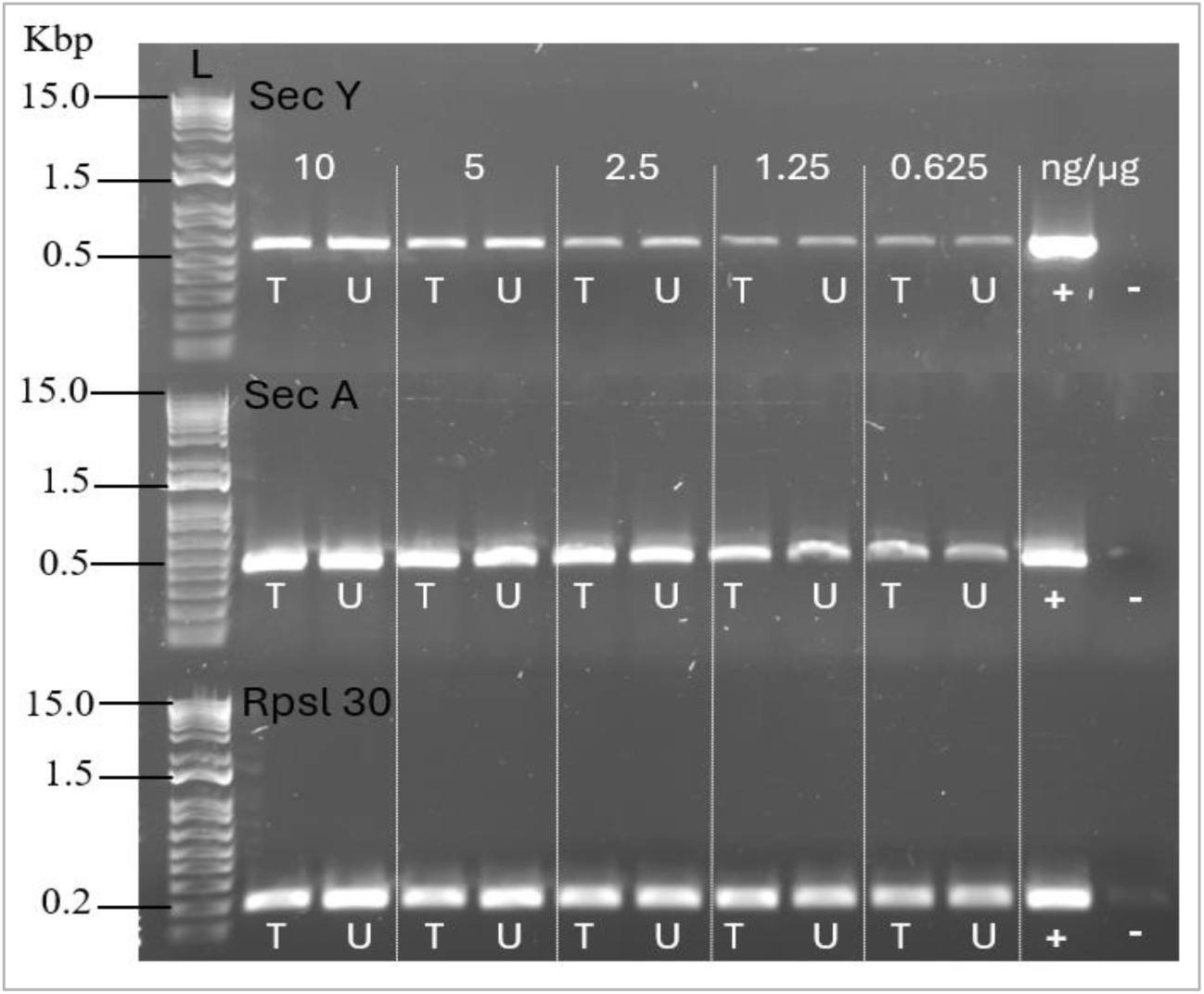
The downregulation of membrane biogenesis is beyond transcription. RT-PCR shows that SecA and SecY, both proteins involved in transmembrane transport, are not downregulated at the RNA level. Treated (T) and untreated (U) samples for each gene were tested at decreasing RNA concentrations (10 – 0.625 ng/µg). Rpsl 30 (200bp), SecY (592bp) and SecA (424bp). (L): Ladder, Invitrogen 1Kb Plus DNA Ladder Cat.No. 10787018.

Importantly, this mode of action is consistent with the absence of membrane disruption observed in the SYTOX Green assay (Figure 4A) and the lack of interaction with bacterial DNA (Figure 4B), supporting a non-lytic, intracellular mechanism. Collectively, these data suggest that H1.2 impairs bacterial growth by disrupting the Sec-dependent protein translocation pathway and subsequent outer membrane assembly, ultimately compromising envelope biogenesis and cellular homeostasis.

### H1.2 forms DNA-based microwebs that entrap bacteria

Given the established role of histones in the formation of neutrophil extracellular traps (NETs), we investigated whether the H1.2 peptide fragment contributes to the formation of NET-like structures and their antibacterial function. To this end, H1.2 was combined with phage DNA and the resulting structures were visualized using scanning electron microscopy (SEM).

SEM analysis revealed that H1.2 readily forms fibrous, web-like structures upon interaction with DNA, resembling synthetic NETs (Figure 6B). These microwebs consisted of DNA fibers decorated with granular H1.2 aggregates, comparable to the architecture observed with full-length calf thymus histones used as a positive control (Figure 6A). However, the H1.2-derived networks appeared less densely packed, likely reflecting the smaller size of the peptide relative to full-length histones.

Importantly, upon exposure to *Pseudomonas aeruginosa*, these H1.2-based microwebs effectively entrapped bacterial cells within the fibrous matrix (Figure 6C), similar to the trapping observed with full-length histone-derived structures (Figure 6D). This physical immobilization suggests that H1.2 contributes to host defense not only through direct antibacterial activity but also by restricting bacterial dissemination.

Collectively, these findings indicate that H1.2 retains the ability to form functional DNA–peptide networks that mimic NET-like architectures, supporting a potential immunomodulatory role in bacterial containment.

## Discussion

Despite increasing recognition of the antimicrobial activity of histones across diverse biological systems, their precise mechanisms of action remain incompletely understood (17). In this study, we identify human histone H1-derived fragments as selective antibacterial effectors and provide mechanistic insight into the activity of the H1.2 peptide against *Pseudomonas aeruginosa*.

We demonstrate that H1-derived peptides exhibit a selective antibacterial profile, with 12 out of 13 fragments significantly inhibiting *P. aeruginosa* growth while showing no activity against other ESKAPE pathogens (Figure 1A). This specificity is maintained across multiple clinical isolates (Figure 1B), pointing to a non-canonical mode of action that does not rely on generalized membrane disruption. Among the tested peptides, H1.2 displayed the most pronounced activity and was therefore selected for detailed characterization. In liquid culture, H1.2 induced rapid bactericidal effects, achieving up to a 5-log reduction in bacterial viability within 2 hours (Figure 1D), although partial regrowth was observed after prolonged incubation. This regrowth likely reflects limited peptide stability or persistence, a common limitation of antimicrobial peptides (36),(37).

Structure–activity relationship analysis revealed that single amino acid substitutions in H1-derived peptides resulted in reduced antibacterial activity (Figure 2A). Notably, these modifications did not significantly alter predicted secondary structures but were associated with a reduction in intramolecular hydrogen bonding (Figure 2B). This suggests that subtle conformational features, rather than overall folding, are critical for maintaining antibacterial function (38),(39). Highlighting the importance of intramolecular interactions in stabilizing bioactive conformations and underscoring the limitations of simplistic sequence-based optimization strategies (40),(41).

Mechanistically, H1.2 does not conform to the classical paradigm of membrane-disrupting antimicrobial peptides. No membrane permeabilization was detected in the SYTOX Green assay (Figure 4A), and electrophoretic mobility shift assays demonstrated that H1.2 does not bind bacterial DNA (Figure 4B). These observations distinguish H1.2 from well-characterized cationic antimicrobial peptides such as LL-37 or defensins, which typically exert their activity through membrane disruption or nucleic acid interaction (42),(43). Instead, our data support a non-lytic and intracellular mechanism of action (44),(45).

Proteomic analysis revealed extensive remodeling of the bacterial proteome following H1.2 treatment, with a pronounced downregulation of proteins involved in outer membrane biogenesis, protein translocation, and folding (Figure 5, Table 4). Specifically, key components of the Sec translocon (SecA, SecD, SecF, SecY, YidC, and YajC), periplasmic chaperones (SurA and Skp), and the β-barrel assembly machinery (BamA, BamB, and Opr86) were significantly reduced. Importantly, RT-PCR analysis demonstrated that the corresponding transcripts, including *secA* and *secY*, were not decreased (Figure 6), indicating that this regulation occurs at a post-transcriptional level. To our knowledge, this represents a previously unrecognized mechanism by which histone-derived peptides interfere with bacterial physiology.

Disruption of these pathways is expected to impair the proper folding, trafficking, and insertion of outer membrane proteins, leading to the accumulation of misfolded proteins and induction of envelope stress (46),(47),(48). Consistent with this, we observed compensatory upregulation of proteins involved in biosynthetic and metabolic processes, including fatty acid synthesis, suggesting activation of adaptive stress responses (49),(50). However, the simultaneous downregulation of quality control proteases, such as MucD, likely exacerbates proteotoxic stress, ultimately compromising bacterial homeostasis and viability (51),(52). While the precise molecular target of H1.2 remains to be identified, one possibility is that the peptide interferes with protein translation, stability, or chaperone-mediated folding, thereby selectively depleting components of membrane biogenesis.

Interestingly, the antibacterial activity of H1.2 is strongly enhanced under acidic conditions (Figure 1C), which are commonly encountered in infected and inflamed tissues (53). In contrast to many antimicrobial peptides, this pH dependency does not correlate with membrane disruption, as H1.2 lacks key protonatable residues typically involved in pH-driven membrane insertion (54),(55). This suggests that acidic environments may instead facilitate peptide uptake, stability, or intracellular targeting, thereby enhancing its antibacterial efficacy in physiologically relevant settings.

In addition to its direct antibacterial effects, H1.2 retains functional properties associated with innate immune defense. We demonstrate that H1.2 forms DNA-based microweb structures resembling neutrophil extracellular traps (NETs) and are capable of entrapping *P. aeruginosa* (Figure 7). Although these structures appear less densely packed than those formed by full-length histones, they effectively immobilize bacterial cells, suggesting a role in limiting bacterial dissemination. This dual functionality—combining intracellular antibacterial activity with extracellular pathogen trapping—highlights the multifaceted role of histone-derived peptides in host defense.

**Figure 7.**
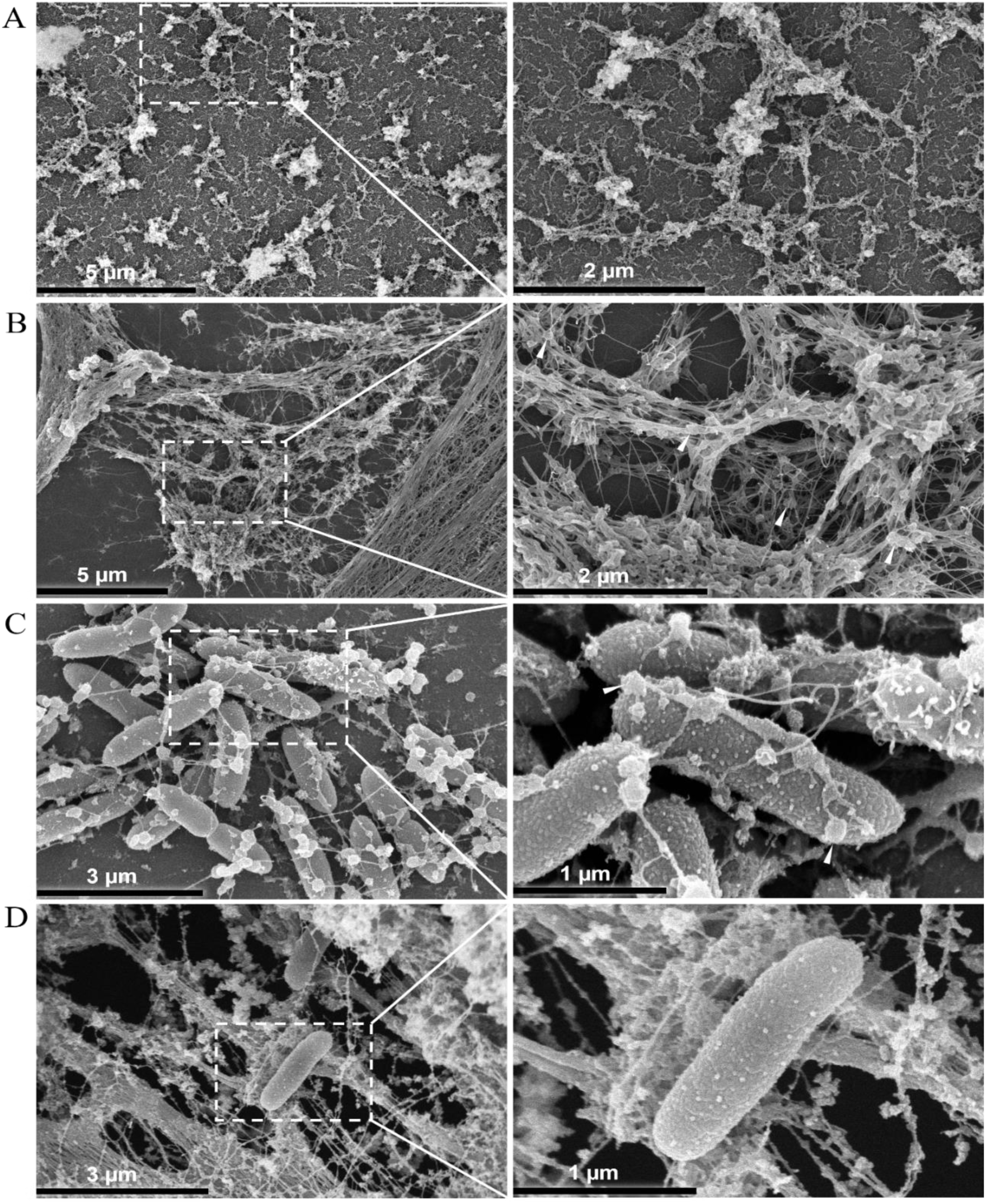
H1.2 peptide forms microwebs that significantly trap bacteria. **(A)** Artificial microwebs formed by phage DNA and full-length histone calf thymus. **(B)** Artificial microwebs formed by phage DNA and H1.2 peptide. **(C)** H1.2 microwebs (DNA + H1.2 + *P. aeruginosa*) trapping bacteria. **(D)** Artificial microwebs (DNA + full length histone + *P. aeruginosa*) trapping bacteria. The white arrows in **(B)** and **(C)** point to the globular H1.2 integrated to DNA fibers.

Importantly, H1.2 exhibited no detectable cytotoxic effects toward THP-1 cells (Figure 3), supporting its selectivity for bacterial targets and reinforcing its potential therapeutic relevance. Moreover, H1.2 did not inhibit biofilm formation (Figure S3), indicating that its activity is primarily directed against planktonic bacteria. This limitation suggests that H1.2 may be most effective during early stages of infection or in combination with agents that disrupt established biofilms.

From a physiological perspective, the identification of H1-derived peptides in human hemofiltrate indicates that these molecules are likely generated in vivo and may function as endogenous antimicrobial effectors. Histones, including linker histone H1, are known to be released into extracellular environments during processes such as cell death, inflammation, and neutrophil extracellular trap formation (56). Proteolytic processing of these proteins in protease-rich environments could give rise to smaller bioactive fragments with enhanced diffusion and specialized antimicrobial functions (16). In this context, H1.2 may represent a naturally occurring effector molecule that contributes to host defense, particularly in acidic and inflammatory microenvironments.

Collectively, our findings support a model in which H1.2 acts as a selective, non-lytic antimicrobial peptide that impairs *P. aeruginosa* growth by disrupting outer membrane biogenesis through post-transcriptional mechanisms. Together with its ability to form NET-like structures and its lack of cytotoxicity, H1.2 represents a promising candidate for the development of novel antimicrobial strategies targeting multidrug-resistant pathogens. Future studies should focus on identifying its direct molecular targets, improving peptide stability, and evaluating its efficacy in in vivo models of infection.

## Credit authorship contribution statement

NJ performed the data investigation assays for the antibacterial activity of H1.2 against *P. aeruginosa*, prepared the data visualization, as well as writing the manuscript. AR identified and predicted H1 peptide fragments and their modifications in silico from the human hemofiltrate library and participated in editing the manuscript. ADS and CC conducted and analyzed the data from the differential proteomics assay, ADS wrote and edited the manuscript, and CC participated in editing the manuscript. CR facilitated the preparation of SEM samples, supervised the capturing of SEM images, and also edited the manuscript. SW and LS were involved in designing experiments and editing. AD supervised the experiments and coordinated the review of the manuscript. JM reviewed and edited the manuscript. BS was responsible for project conceptualization, supervision of the experiments, and editing the manuscript. All authors contributed to the article and approved the submitted version.

## Declaration of generative AI use

During the preparation of this work, the authors used ChatGPT 5.2 architecture by OpenAI and *Perplexity AI* (https://www.perplexity.ai), powered by GPT-5, OpenAI, in order to assist with literature review, experimental troubleshooting, data interpretation, and manuscript text. The AI tools were used to refine scientific writing and improve the clarity of presentation. After using these tools, the authors reviewed and edited the content as needed and take full responsibility for the content of the published article.

## Acknowledgements

NJ is part of, and would like to acknowledge and thank, the International Graduate School in Molecular Medicine Ulm (IGradU).

## References

1. Murray CJL, Ikuta KS, Sharara F, Swetschinski L, Robles Aguilar G, Gray A, et al. Global burden of bacterial antimicrobial resistance in 2019: a systematic analysis. The Lancet. 2022;399(10325):629–55.

2. Ferraz MP. Antimicrobial Resistance: The Impact from and on Society According to One Health Approach. Societies. 2024;14(9):187.

3. Mukhopadhyay S, Bharath Prasad AS, Mehta CH, Nayak UY. Antimicrobial peptide polymers: no escape to ESKAPE pathogens-a review. World J Microbiol Biotechnol. 2020;36(9):131.

4. Hancock RE, Patrzykat A. Clinical development of cationic antimicrobial peptides: from natural to novel antibiotics. Curr Drug Targets Infect Disord. 2002;2(1):79–83.

5. Hancock REW, Lehrer R. Cationic peptides: a new source of antibiotics. Trends in Biotechnology. 1998;16(2):82–8.

6. Scott MG, Yan H, Hancock RE. Biological properties of structurally related alpha-helical cationic antimicrobial peptides. Infect Immun. 1999;67(4):2005–9.

7. Hancock RE, Sahl HG. Antimicrobial and host-defense peptides as new anti-infective therapeutic strategies. Nat Biotechnol. 2006;24(12):1551–7.

8. Kamaruzzaman NF, Tan LP, Hamdan RH, Choong SS, Wong WK, Gibson AJ, et al. Antimicrobial Polymers: The Potential Replacement of Existing Antibiotics? Int J Mol Sci. 2019;20(11).

9. Kumar P, Kizhakkedathu JN, Straus SK. Antimicrobial Peptides: Diversity, Mechanism of Action and Strategies to Improve the Activity and Biocompatibility In Vivo. Biomolecules. 2018;8(1).

10. Hsu CH, Chen C, Jou ML, Lee AY, Lin YC, Yu YP, et al. Structural and DNA-binding studies on the bovine antimicrobial peptide, indolicidin: evidence for multiple conformations involved in binding to membranes and DNA. Nucleic Acids Res. 2005;33(13):4053–64.

11. Jenssen H, Hamill P, Hancock RE. Peptide antimicrobial agents. Clin Microbiol Rev. 2006;19(3):491–511.

12. Mookherjee N, Anderson MA, Haagsman HP, Davidson DJ. Antimicrobial host defence peptides: functions and clinical potential. Nature Reviews Drug Discovery. 2020;19(5):311–32.

13. Wang G. Human Antimicrobial Peptides and Proteins. Pharmaceuticals. 2014;7(5):545–94.

14. Lai Y, Gallo RL. AMPed up immunity: how antimicrobial peptides have multiple roles in immune defense. Trends Immunol. 2009;30(3):131–41.

15. Miller BF, Abrams R, Dorfman A, Klein M. Antibacterial Properties of Protamine and Histone. Science. 1942;96(2497):428–30.

16. Li X, Ye Y, Peng K, Zeng Z, Chen L, Zeng Y. Histones: The critical players in innate immunity. Front Immunol. 2022;13:1030610.

17. Rose FR, Bailey K, Keyte JW, Chan WC, Greenwood D, Mahida YR. Potential role of epithelial cell-derived histone H1 proteins in innate antimicrobial defense in the human gastrointestinal tract. Infect Immun. 1998;66(7):3255–63.

18. Vorobjeva NV, Chernyak BV. NETosis: Molecular Mechanisms, Role in Physiology and Pathology. Biochemistry (Mosc). 2020;85(10):1178–90.

19. Wang Y, Du C, Zhang Y, Zhu L. Composition and Function of Neutrophil Extracellular Traps. Biomolecules. 2024;14(4).

20. Urban CF, Ermert D, Schmid M, Abu-Abed U, Goosmann C, Nacken W, et al. Neutrophil Extracellular Traps Contain Calprotectin, a Cytosolic Protein Complex Involved in Host Defense against Candida albicans. PLOS Pathogens. 2009;5(10):e1000639.

21. Brinkmann V, Reichard U, Goosmann C, Fauler B, Uhlemann Y, Weiss DS, et al. Neutrophil extracellular traps kill bacteria. science. 2004;303(5663):1532–5.

22. Song Y, Kadiyala U, Weerappuli P, Valdez JJ, Yalavarthi S, Louttit C, et al. Antimicrobial Microwebs of DNA-Histone Inspired from Neutrophil Extracellular Traps. Adv Mater. 2019;31(14):e1807436.

23. Yang T, Yu J, Ahmed T, Nguyen K, Nie F, Zan R, et al. Synthetic neutrophil extracellular traps dissect bactericidal contribution of NETs under regulation of α-1-antitrypsin. Sci Adv. 2023;9(17):eadf2445.

24. Kim HS, Cho JH, Park HW, Yoon H, Kim MS, Kim SC. Endotoxin-neutralizing antimicrobial proteins of the human placenta. J Immunol. 2002;168(5):2356–64.

25. Bolton SJ, Perry VH. Histone H1; a neuronal protein that binds bacterial lipopolysaccharide. J Neurocytol. 1997;26(12):823–31.

26. Lee DY, Huang CM, Nakatsuji T, Thiboutot D, Kang SA, Monestier M, et al. Histone H4 is a major component of the antimicrobial action of human sebocytes. J Invest Dermatol. 2009;129(10):2489–96.

27. Tanaka Y, Yamanaka N, Koyano I, Hasunuma I, Kobayashi T, Kikuyama S, et al. Dual Roles of Extracellular Histone H3 in Host Defense: Its Differential Regions Responsible for Antimicrobial and Cytotoxic Properties and Their Modes of Action. Antibiotics (Basel). 2022;11(9).

28. Yan J, Wang K, Dang W, Chen R, Xie J, Zhang B, et al. Two hits are better than one: membrane-active and DNA binding-related double-action mechanism of NK-18, a novel antimicrobial peptide derived from mammalian NK-lysin. Antimicrob Agents Chemother. 2013;57(1):220–8.

29. Cox J, Neuhauser N, Michalski A, Scheltema RA, Olsen JV, Mann M. Andromeda: a peptide search engine integrated into the MaxQuant environment. J Proteome Res. 2011;10(4):1794–805.

30. Canè C, Casciaro B, Di Somma A, Loffredo MR, Puglisi E, Battaglia G, et al. The antimicrobial peptide Esc(1-21)-1c increases susceptibility of Pseudomonas aeruginosa to conventional antibiotics by decreasing the expression of the MexAB-OprM efflux pump. Frontiers in Chemistry. 2023;Volume 11 - 2023.

31. Schütz D, Rode S, Read C, Müller JA, Glocker B, Sparrer KMJ, et al. Viral Transduction Enhancing Effect of EF-C Peptide Nanofibrils Is Mediated by Cellular Protrusions. Advanced Functional Materials. 2021;31(40):2104814.

32. Waghu FH, Barai RS, Gurung P, Idicula-Thomas S. CAMPR3: a database on sequences, structures and signatures of antimicrobial peptides. Nucleic Acids Res. 2016;44(D1):D1094–7.

33. Meher PK, Sahu TK, Saini V, Rao AR. Predicting antimicrobial peptides with improved accuracy by incorporating the compositional, physico-chemical and structural features into Chou’s general PseAAC. Sci Rep. 2017;7:42362.

34. Lin TT, Yang LY, Lu IH, Cheng WC, Hsu ZR, Chen SH, et al. AI4AMP: an Antimicrobial Peptide Predictor Using Physicochemical Property-Based Encoding Method and Deep Learning. mSystems. 2021;6(6):e0029921.

35. Starr TN, Thornton JW. Exploring protein sequence-function landscapes. Nat Biotechnol. 2017;35(2):125–6.

36. Ma X, Aminov R, Franco OL, de la Fuente-Nunez C, Wang G, Wang J. Editorial: Antimicrobial peptides and their druggability, bio-safety, stability, and resistance. Frontiers in Microbiology. 2024;Volume 15 - 2024.

37. Lai Z, Yuan X, Chen H, Zhu Y, Dong N, Shan A. Strategies employed in the design of antimicrobial peptides with enhanced proteolytic stability. Biotechnology Advances. 2022;59:107962.

38. Mitra M, Asad M, Kumar S, Yadav K, Chaudhary S, Bhavesh NS, et al. Distinct Intramolecular Hydrogen Bonding Dictates Antimicrobial Action of Membrane-Targeting Amphiphiles. The Journal of Physical Chemistry Letters. 2019;10(4):754–60.

39. Sakamoto T, Koga Y, Hikota M, Matsuki K, Murakami M, Kikkawa K, et al. Design and synthesis of novel 5-(3,4,5-trimethoxybenzoyl)-4-aminopyrimidine derivatives as potent and selective phosphodiesterase 5 inhibitors: Scaffold hopping using a pseudo-ring by intramolecular hydrogen bond formation. Bioorganic & Medicinal Chemistry Letters. 2014;24(22):5175–80.

40. Giordanetto F, Tyrchan C, Ulander J. Intramolecular Hydrogen Bond Expectations in Medicinal Chemistry. ACS Medicinal Chemistry Letters. 2017;8(2):139–42.

41. Ahmed YM, Mohamed GG. Synthesis, spectral characterization, antimicrobial evaluation and molecular docking studies on new metal complexes of novel Schiff base derived from 4,6-dihydroxy-1,3-phenylenediethanone. Journal of Molecular Structure. 2022;1256:132496.

42. Benfield AH, Henriques ST. Mode-of-Action of Antimicrobial Peptides: Membrane Disruption vs. Intracellular Mechanisms. Front Med Technol. 2020;2:610997.

43. Pachón-Ibáñez ME, Smani Y, Pachón J, Sánchez-Céspedes J. Perspectives for clinical use of engineered human host defense antimicrobial peptides. FEMS Microbiology Reviews. 2017;41(3):323–42.

44. Nicolas P. Multifunctional host defense peptides: intracellular-targeting antimicrobial peptides. The FEBS Journal. 2009;276(22):6483–96.

45. Lemaire S, Trinh T-T, Le H-T, Tang S-C, Hincke M, Wellman-Labadie O, et al. Antimicrobial effects of H4-(86–100), histogranin and related compounds – possible involvement of DNA gyrase. The FEBS Journal. 2008;275(21):5286–97.

46. Alvira S, Watkins DW, Troman LA, Allen WJ, Lorriman JS, Degliesposti G, et al. Inter-membrane association of the Sec and BAM translocons for bacterial outer-membrane biogenesis. eLife. 2020;9:e60669.

47. Mitchell AM, Silhavy TJ. Envelope stress responses: balancing damage repair and toxicity. Nat Rev Microbiol. 2019;17(7):417–28.

48. Ranava D, Yang Y, Orenday-Tapia L, Rousset F, Turlan C, Morales V, et al. Lipoprotein DolP supports proper folding of BamA in the bacterial outer membrane promoting fitness upon envelope stress. eLife. 2021;10:e67817.

49. Xue J, Li S, Wang L, Zhao Y, Zhang L, Zheng Y, et al. Enhanced fatty acid biosynthesis by Sigma28 in stringent responses contributes to multidrug resistance and biofilm formation in Helicobacter pylori. Antimicrob Agents Chemother. 2024;68(9):e0085024.

50. Bernal P, Segura A, Ramos J-L. Compensatory role of the cis-trans-isomerase and cardiolipin synthase in the membrane fluidity of Pseudomonas putida DOT-T1E. Environmental Microbiology. 2007;9(7):1658–64.

51. Yorgey P, Rahme LG, Tan M-W, Ausubel FM. The roles of mucD and alginate in the virulence of Pseudomonas aeruginosa in plants, nematodes and mice. Molecular Microbiology. 2001;41(5):1063–76.

52. Pan KL, Hsiao HC, Weng CL, Wu MS, Chou CP. Roles of DegP in prevention of protein misfolding in the periplasm upon overexpression of penicillin acylase in Escherichia coli. J Bacteriol. 2003;185(10):3020–30.

53. Hajjar S, Zhou X. pH sensing at the intersection of tissue homeostasis and inflammation. Trends Immunol. 2023;44(10):807–25.

54. Hitchner MA, Santiago-Ortiz LE, Necelis MR, Shirley DJ, Palmer TJ, Tarnawsky KE, et al. Activity and characterization of a pH-sensitive antimicrobial peptide. Biochim Biophys Acta Biomembr. 2019;1861(10):182984.

55. Malik E, Dennison SR, Harris F, Phoenix DA. pH Dependent Antimicrobial Peptides and Proteins, Their Mechanisms of Action and Potential as Therapeutic Agents. Pharmaceuticals (Basel). 2016;9(4).

56. Chen R, Kang R, Fan XG, Tang D. Release and activity of histone in diseases. Cell Death & Disease. 2014;5(8):e1370–e.

