## Supplemental Figures for "Human Histone Fragments Display Antibacterial Properties against *Pseudomonas aeruginosa*"

### Supplementary Figures

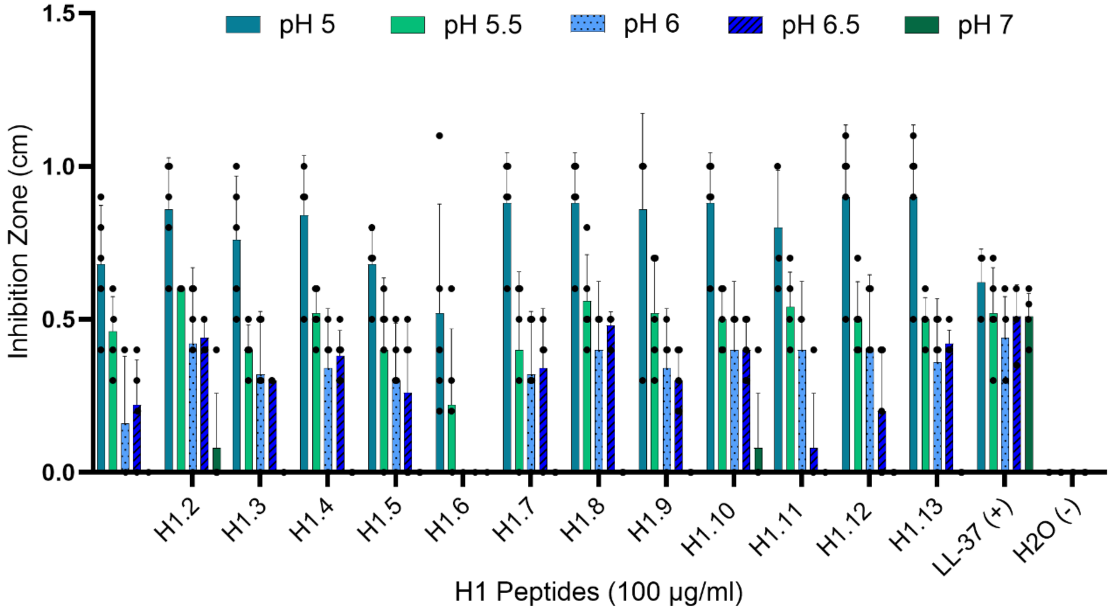


were tested against *P. aeruginosa* in different pH media in a radial diffusion assay.

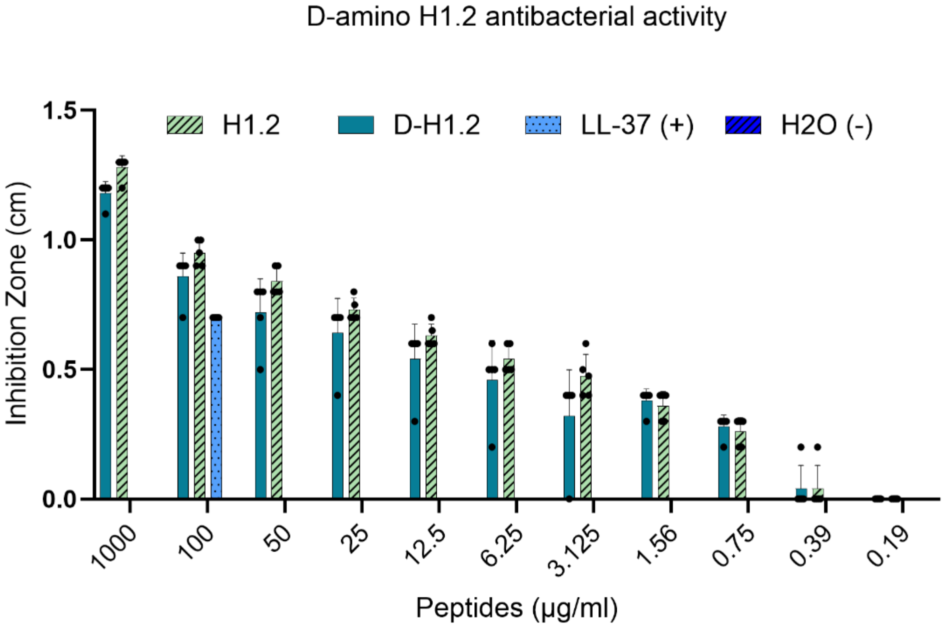


**Figure S2**. **H1.2 and D-amino H1.2 peptides in RDA.** The antibacterial activity tested at lower concentrations of H1.2 and D-amino H1.2 against *P. aeruginosa* in RDA assay. Both forms of the peptide the L-amino acid and the D-amino acid function similarly in inhibiting the growth of *P. aeruginosa*.

**A.**

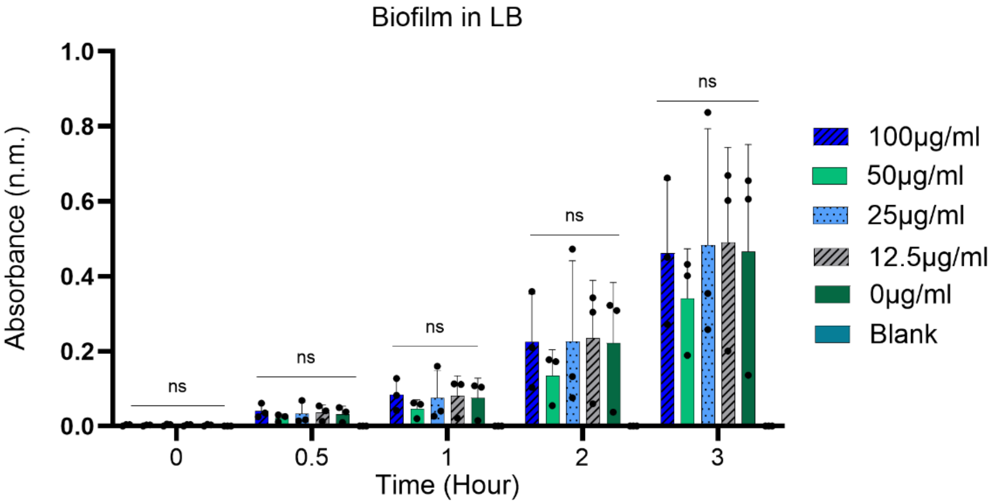


**B.**

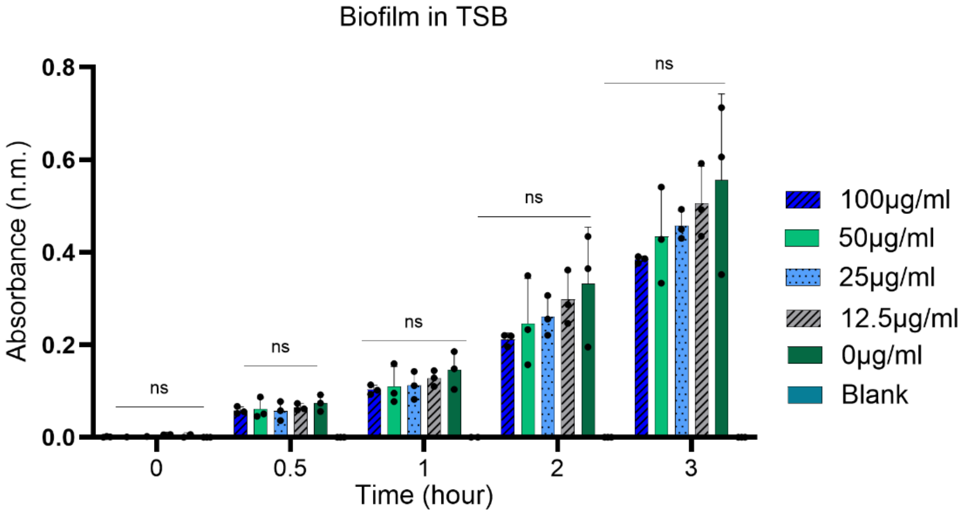


**C.**

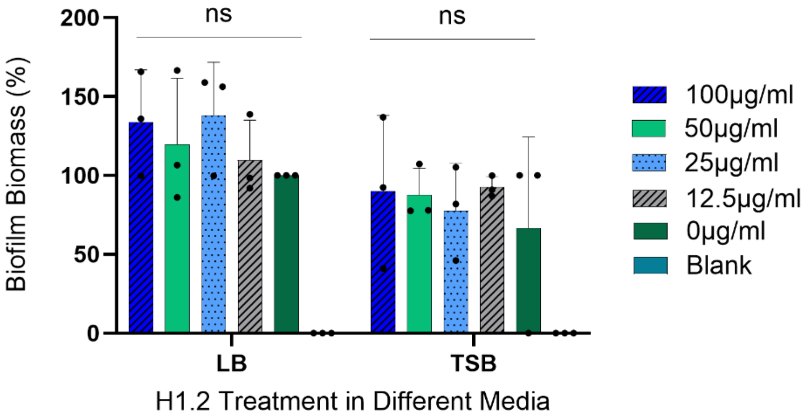


**Figure S3**. **H1.2 Histone peptide does not inhibit biofilm formation.** Incubating H1.2 of different concentrations with *P. aeruginosa* overnight at 37°C does not interfere with the viability of the bacteria within the formed biofilm in LB **(A)** or TSB **(B)** media. **(C).** Crystal violet assay of H1.2 tested against *P. aeruginosa* biofilm in LB and TSB media shows no significant reduction in the biofilm biomass formation.
