## Supplemental Tables for "Human Histone Fragments Display Antibacterial Properties against *Pseudomonas aeruginosa*"

### Supplementary Tables

**Supplementary Tables. Detailed analysis of proteomics data.** MaxQuant analysis using UniProt *P. aeruginosa* shows some up-regulated **(Table 1)** and down-regulated **(Table 2)** proteins after treatment of bacteria with H1.2 peptide. The tables show the differentially expressed proteins, including their UniProt IDs, gene names, biological function, and fold changes (FC).

#### Table 1: Up-regulated Proteins

| Protein ID | Protein name | Gene | Biological function | FC |
|---|---|---|---|---|
| O54439 | Acyl carrier protein 1 | *acpP1* | fatty acid biosynthesis | 4,946 |
| Q9HTX6 | Glycine cleavage system H protein 2 | *gcvH2* | glycine decarboxylation | 3,153 |
| Q9HTQ4 | Cytochrome c5 | *cycB* | electron transport | 3,042 |
| P11724 | Ornithine carbamoyltransferase, anabolic | *argF* | amino-acid biosynthesis | 2,536 |
| Q9HW32 | Insulin-cleaving metalloproteinase outer membrane protein | *icmP* | metal ion binding (mol) | 2,350 |
| E1JGJ8 | Peptide chain release factor 2 | *prfB* | protein biosynthesis | 2,266 |
| Q9X2T1 | Thioredoxin | *trxA* | electron transport | 2,092 |
| Q9HV46 | Transcription elongation factor GreA | *greA* | transcription | 1,906 |
| Q9HWD5 | Large ribosomal subunit protein uL3 | *rplC* | translation | 1,895 |
| P00282 | Azurin | *azu* | electron transport | 1,772 |
| Q9I0A0 | Translation initiation factor IF-3 | *infC* | protein biosynthesis | 1,629 |
| Q9HWF4 | Large ribosomal subunit protein uL15 | *rplO* | translation | 1,611 |
| Q9HUN2 | Large ribosomal subunit protein bL9 | *rplI* | translation | 1,444 |
| O82853 | Ribosome-recycling factor | *frr* | protein biosynthesis | 1,423 |

| | | | | |
|---|---|---|---|---|
| Q9I2U9 | Peptidyl-prolyl cis-trans isomerase | *ppiB* | protein folding | 1,412 |
| P95459 | Major cold shock protein CspA | *cspA* | stress response | 1,410 |
| P13982 | Carbamate kinase | *arcC* | arginine metabolism | 1,402 |
| O82851 | Elongation factor Ts | *tsf* | protein biosynthesis | 1,393 |
| P57668 | Thiol peroxidase | *tpx* | cellular response | 1,387 |
| Q9HZC7 | DUF1329 domain-containing protein | | unknown | 1,381 |
| Q9HUD0 | Large ribosomal subunit protein bL31 | *rpmE* | translation | 1,380 |
| Q9HV60 | BON domain-containing protein | | response to osmotic stress | 1,360 |
| P29363 | Threonine synthase | *thrC* | amino-acid biosynthesis | 1,353 |
| Q9HWE4 | Small ribosomal subunit protein uS17 | *rpsQ* | translation | 1,341 |
| Q9HU11 | Peptidoglycan-binding protein LysM | | response to potassium ion | 1,326 |
| Q9HWC7 | Large ribosomal subunit protein uL10 | *rplJ* | translation | 1,325 |
| Q9HVI7 | Serine hydroxymethyltransferase 3 | *glyA3* | amino-acid biosynthesis | 1,309 |
| Q9I3B2 | 3-oxoacyl-[acyl-carrier-protein] synthase 1 | *fabB* | fatty acid biosynthesis | 1,295 |
| Q9HUM2 | Protein HflK | *hflK* | proteolysis | 1,280 |
| Q9HXU9 | DUF4398 domain-containing protein | | unknown | 1,278 |
| Q9HTR6 | Nitrogen regulatory protein P-II 2 | *glnK* | cellular response to antibiotic | 1,237 |
| Q9HZP7 | Electron transfer flavoprotein subunit alpha | *etfA* | electron transport | 1,219 |
| Q9I5E3 | Citrate synthase | *prpC* | carbohydrate metabolic process | 1,209 |

### Table 2: Down-regulated Proteins

| Protein IDs | Protein name | Gene | Biological function | FC |
|---|---|---|---|---|
| Q9I5A7 | OmpA-like domain-containing protein | | monoatomic ion transport | 0,800 |
| Q9HVV7 | Lipopolysaccharide export system protein LptH | *lptH* | lipopolysaccharide tranposrt | 0,798 |
| Q9HXV4 | Adenylate kinase | *adk* | nucleotide biosynthesis | 0,797 |
| Q9HT20 | ATP synthase subunit beta | *atpD* | atp synthesis | 0,795 |
| Q9I671 | glutaryl-CoA dehydrogenase (ETF) | *gcdH* | biosynthetic process | 0,794 |
| P09591 | Elongation factor Tu | *tufA* | translation elongation | 0,789 |
| Q9HVY4 | Ubiquinol-cytochrome c reductase iron-sulfur subunit | | oxidoreductase activity | 0,787 |
| Q9I685 | Adenosylhomocysteinase | *ahcY* | metabolic process | 0,784 |
| Q9I2V5 | Aconitate hydratase B | *acnB* | tricarboxylic acid cycle | 0,781 |
| P53641 | Superoxide dismutase [Fe] | *sodB* | superoxide dismutase activity | 0,777 |
| Q9HVA2 | Ketol-acid reductoisomerase (NADP(+)) | *ilvC* | biosynthetic process | 0,776 |
| Q9HVT7 | Aspartyl/glutamyl-tRNA(Asn/Gln) amidotransferase subunit B | *gatB* | translation | 0,772 |
| Q9I0L4 | Isocitrate dehydrogenase [NADP] | *idh* | oxidoreductase (mol) | 0,771 |
| Q9HWD1 | Small ribosomal subunit protein uS7 | *rpsG* | translation | 0,765 |
| Q9HTN8 | Large ribosomal subunit protein bL28 | *rpmB* | translation | 0,764 |
| Q9HVY3 | Small ribosomal subunit protein uS9 | *rpsI* | translation | 0,764 |
| Q9HYK7 | ferredoxin--NADP(+) reductase | *fpr* | catabolic process | 0,763 |
| O30508 | Succinylornithine transaminase/acetylornithine aminotransferase | *aruC* | biosynthetic process | 0,758 |

| | | | | |
|---|---|---|---|---|
| Q9HTZ7 | Phosphoenolpyruvate carboxykinase | *pckA* | gluconeogenesis | 0,758 |
| Q9HTL9 | RidA family protein | | catabolic process | 0,756 |
| G3XD11 | PhoP/Q and low Mg2+ inducible outer membrane protein H1 | *oprH* | lipopolysaccharide binding | 0,752 |
| Q9I4I1 | Ribonucleoside-diphosphate reductase | *nrdA* | deoxyribonucleotide synthesis | 0,751 |
| Q9HXM5 | Inosine-5'-monophosphate dehydrogenase | *guaB* | biosynthetic process | 0,750 |
| Q9HTN2 | Acetylglutamate kinase | *argB* | biosynthetic process | 0,749 |
| Q9HWC9 | DNA-directed RNA polymerase subunit beta' | *rpoC* | dna-templated transcription | 0,749 |
| Q9HT18 | ATP synthase subunit alpha | *atpA* | atp synthesis | 0,748 |
| Q9I3G1 | Cytochrome c oxidase, cbb3-type, CcoO subunit | *ccoO2* | cytochrome-c oxidase activity | 0,748 |
| Q9I4S1 | Glycine zipper 2TM domain-containing protein | | | 0,746 |
| Q51487 | Outer membrane protein OprM | *oprM* | response to antibiotic/ transmembrane transport | 0,743 |
| Q9HUM5 | ATP phosphoribosyltransferase regulatory subunit | *hisZ* | biosynthetic process | 0,743 |
| Q9HV42 | Protein GrpE | *grpE* | protein folding | 0,741 |
| Q9I0L5 | Isocitrate dehydrogenase [NADP] | *icd* | oxidoreductase (mol) | 0,741 |
| Q9HWE7 | Large ribosomal subunit protein uL5 | *rplE* | translation | 0,740 |
| Q9I2A6 | 3-hydroxybutyrate dehydrogenase | *bdhA* | oxidoreductase (mol) | 0,739 |
| Q9I420 | Carboxypeptidase regulatory-like domain-containing protein | | | 0,737 |
| P08308 | Ornithine carbamoyltransferase | *arcB* | biosynthetic process | 0,732 |
| Q9HZM8 | Ribonuclease E | *rne* | trna processing | 0,732 |
| Q9HUM9 | Small ribosomal subunit protein bS6 | *rpsF* | translation | 0,731 |
| G3XD30 | Type IV pilus inner membrane component PilN | *pilN* | type iv pilus assembly | 0,729 |
| Q51567 | Succinate--CoA ligase | *sucD* | biosynthetic process | 0,728 |
| Q9I5H0 | Folate-binding protein | | iron-sulfur cluster assembly | 0,727 |

| | | | | |
|---|---|---|---|---|
| P40947 | Single-stranded DNA-binding protein | *ssb* | dna binding | 0,723 |
| Q9HVC5 | Ribose-phosphate pyrophosphokinase | *prs* | biosynthetic process | 0,723 |
| Q9I6Z3 | Alkyl hydroperoxide reductase C | *ahpC* | cellular response to stress | 0,720 |
| Q9HWE0 | Large ribosomal subunit protein uL22 | *rplV* | translation | 0,718 |
| Q59636 | Nucleoside diphosphate kinase | *ndk* | biosynthetic process | 0,717 |
| Q9HVL6 | Large ribosomal subunit protein bL21 | *rplU* | translation | 0,717 |
| Q9HVN5 | Chaperone protein ClpB | *clpB* | protein refolding | 0,716 |
| O52762 | Catalase | *katA* | catabolic process | 0,714 |
| P37798 | Biotin carboxylase | *accC* | biosynthetic process | 0,713 |
| Q9I5Q3 | Tyrosine--tRNA ligase 2 | *tyrS2* | protein biosynthesis | 0,712 |
| Q9ZN70 | Exopolyphosphatase | *ppx* | biosynthetic process | 0,712 |
| O52759 | Small ribosomal subunit protein uS4 | *rpsD* | translation | 0,709 |
| Q9I687 | Methylenetetrahydrofolate reductase | *metF* | amino-acid biosynthesis | 0,703 |
| Q9HWW1 | Outer membrane protein OprG | *oprG* | protein transport | 0,703 |
| Q59637 | Pyruvate dehydrogenase E1 component | *aceE* | pyruvate dehydrogenase activity | 0,700 |
| Q9HWD8 | Large ribosomal subunit protein uL2 | *rplB* | cytoplasmic translation | 0,699 |
| Q9HWD6 | Large ribosomal subunit protein uL4 | *rplD* | translation | 0,696 |
| O50273 | Sulfate adenylyltransferase subunit 2 | *cysD* | biosynthetic process | 0,694 |
| Q9HXP3 | Formate-dependent phosphoribosylglycinamide formyltransferase | *purT* | purine biosynthesis | 0,692 |
| Q9I5Y4 | Phosphoglycerate kinase | *pgk* | glycolysis | 0,687 |
| Q9HTP2 | Probable aldehyde dehydrogenase | | oxidoreductase activity | 0,686 |
| P13981 | Arginine deiminase | *arcA* | catabolic process | 0,685 |

| | | | | |
|---|---|---|---|---|
| Q9I0K4 | Isocitrate lyase | | biofilm formation | 0,679 |
| Q9HWD9 | Small ribosomal subunit protein uS19 | *rpsS* | translation | 0,675 |
| Q9HVD1 | Lipid A deacylase PagL | *pagL* | metabolic process | 0,671 |
| Q9I3D1 | Dihydrolipoyl dehydrogenase | *lpdG* | oxidoreductase (mol) | 0,670 |
| Q9HVY2 | Large ribosomal subunit protein uL13 | *rplM* | translation | 0,669 |
| Q9HVX1 | Probable periplasmic serine endoprotease DegP-like | *algW* | proteolysis | 0,668 |
| Q9HWX4 | 3,4-dihydroxy-2-butanone 4-phosphate synthase | *ribB* | riboflavin biosynthesis | 0,665 |
| Q9HWF7 | Small ribosomal subunit protein uS13 | *rpsM* | translation | 0,664 |
| Q9HWD4 | Small ribosomal subunit protein uS10 | *rpsJ* | translation | 0,663 |
| Q9I765 | oligopeptidase A | *prlC* | metabolic process | 0,663 |
| Q9HTD9 | alcohol dehydrogenase | *adhA* | zinc ion binding | 0,662 |
| Q9I3G2 | Cbb3-type cytochrome c oxidase subunit | *ccoP2* | electron transport | 0,660 |
| P38098 | Carbamoyl phosphate synthase small chain | *carA* | biosynthetic process | 0,656 |
| Q9X6W6 | Bifunctional protein PyrR | *pyrR* | transcription | 0,656 |
| Q59643 | Delta-aminolevulinic acid dehydratase | *hemB* | biosynthetic process | 0,653 |
| G3XD89 | Copper transport outer membrane porin OprC | *oprC* | transmembrane transport | 0,652 |
| Q9I3C5 | Chaperone protein HtpG | *htpG* | stress response | 0,650 |
| Q9I5Z0 | S-adenosylmethionine synthase | *metK* | metabolism | 0,649 |
| Q9HU65 | Glutamine synthetase | *glnA* | biosynthetic process | 0,648 |
| Q9I574 | Membrane-bound lysozyme inhibitor of C-type lysozyme | *mliC* | | 0,648 |
| Q9I4W3 | 4-hydroxy-tetrahydrodipicolinate synthase | *dapA* | amino-acid biosynthesis | 0,646 |
| Q9HVS6 | DUF541 domain-containing protein | | dna damage response | 0,645 |
| Q9HXU0 | Lysine--tRNA ligase | *lysS* | protein biosynthesis | 0,645 |

| | | | | |
|---|---|---|---|---|
| Q9HVW4 | Toluene tolerance protein | | | 0,644 |
| Q9HTD0 | Biotin carboxylase | | biosynthetic process | 0,643 |
| Q9I5F6 | Bifunctional protein PutA | *putA* | metabolism | 0,643 |
| Q9I5U9 | PrkA AAA domain-containing protein | | protein kinase activity (mol) | 0,643 |
| Q9HUA7 | Probable binding protein component of ABC transporter | | | 0,637 |
| Q9I0X8 | DUF2185 domain-containing protein | | unknown | 0,637 |
| Q9HU73 | Fructose-1,6-bisphosphatase class 1 | *fbp* | metabolic process | 0,629 |
| Q9HTQ1 | RidA family protein | | no information | 0,628 |
| Q9HZJ5 | DNA topoisomerase 1 | *topA* | dna-binding (mol) | 0,626 |
| O50274 | Bifunctional enzyme CysN/CysC | *cysNC* | metabolic process | 0,625 |
| Q9HWE1 | Small ribosomal subunit protein uS3 | *rpsC* | translation | 0,624 |
| Q9HYY3 | Periplasmic tail-specific protease | *prc* | proteolysis | 0,622 |
| Q9HTD1 | Probable transcarboxylase subunit | | ion transport | 0,621 |
| Q9I0M4 | Outer-membrane lipoprotein carrier protein | *lolA* | protein transport | 0,620 |
| Q9I1R3 | Probable binding protein component of ABC transporter | | ion transport (mol) | 0,618 |
| Q9I405 | Amino acid ABC transporter ATP binding protein | | transport | 0,617 |
| Q9HZ55 | Ribosomal large subunit pseudouridine synthase B | *rluB* | translation | 0,616 |
| Q9HZ48 | Probable binding protein component of ABC sugar transporter | | protein transport | 0,614 |
| Q9I3F5 | Aconitate hydratase A | *acnA* | tricarboxylic acid cycle-electron transport | 0,612 |
| Q9I4I2 | Ribonucleoside-diphosphate reductase subunit beta | *nrdB* | deoxyribonucleotide synthesis | 0,612 |
| Q9HVY6 | Probable cytochrome c1 | | metal ion binding | 0,607 |
| Q9I3D5 | Succinate dehydrogenase flavoprotein subunit | *sdhA* | electron transport | 0,606 |
| Q9I0J5 | NADH-quinone oxidoreductase subunit H | *nuoH* | aerobic respiration | 0,604 |

| | | | | |
|---|---|---|---|---|
| Q9I0H4 | Flavohemoprotein | *hmp* | cellular response to stress | 0,602 |
| G3XD51 | Type IV pilus inner membrane component PilO | *pilO* | type iv pilus assembly | 0,601 |
| Q9HX12 | MaoC-like domain-containing protein | | | 0,601 |
| Q9HYU8 | Probable HIT family protein | | metabolic process | 0,601 |
| Q9HUC0 | Ubiquinone/menaquinone biosynthesis C-methyltransferase UbiE | *ubiE* | biosynthetic process | 0,600 |
| Q9HXY5 | Skp-like protein | | protein folding | 0,600 |
| Q9HYC7 | Methionine--tRNA ligase | *metG* | protein biosynthesis | 0,600 |
| Q03456 | Ferric uptake regulation protein | *fur* | dna binding | 0,597 |
| Q9HYQ8 | ADP-L-glycero-D-manno-heptose-6-epimerase | *hldD* | biosynthetic process | 0,597 |
| Q9HY13 | Xylulose 5-phosphate/Fructose 6-phosphate phosphoketolase N-terminal domain-containing protein | | metabolic process | 0,597 |
| Q9HWV5 | Probable aldehyde dehydrogenase | | phenylacetaldehyde dehydrogenase activity (mol) | 0,595 |
| Q9I3D3 | oxoglutarate dehydrogenase (succinyl-transferring) | *sucA* | tricarboxylic acid cycle-electron transport | 0,594 |
| Q9HV67 | Glucose-6-phosphate isomerase | *pgi* | metabolic process | 0,593 |
| Q9I4K8 | Probable oxidoreductase | | oxidoreductase (mol) | 0,593 |
| Q9HZK4 | Glyceraldehyde-3-phosphate dehydrogenase | | metabolic process | 0,593 |
| P27726 | Glyceraldehyde-3-phosphate dehydrogenase | *gap* | metabolic process | 0,590 |
| Q9I5Y1 | Fructose-bisphosphate aldolase | *fba* | glycolysis | 0,589 |
| Q9HTQ6 | Xanthine phosphoribosyltransferase | *xpt* | metabolic process | 0,586 |
| Q9HVM4 | Isoleucine--tRNA ligase | *ileS* | isoleucyl-trnaaminoacylation | 0,586 |
| Q9I0A2 | Large ribosomal subunit protein bL20 | *rplT* | translation | 0,585 |
| P43335 | Pterin-4-alpha-carbinolamine dehydratase | *phhB* | biosynthetic process | 0,583 |
| Q9HVA1 | Acetolactate synthase small subunit | *ilvH* | biosynthetic process | 0,580 |

| | | | | |
|---|---|---|---|---|
| P43336 | Aromatic-amino-acid aminotransferase | *phhC* | catabolic process | 0,579 |
| Q9I6J6 | Probable periplasmic polyamine binding protein | | transport | 0,579 |
| Q9I047 | Transaldolase | *tal* | metabolic process | 0,579 |
| Q9I636 | Malate synthase G | *glcB* | catabolic process | 0,578 |
| Q9HZ98 | Heat-shock protein IbpA | *ibpA* | stress response | 0,576 |
| Q9HVL9 | Glutamate 5-kinase | *proB* | biosynthetic process | 0,574 |
| Q9I4W5 | thioredoxin-dependent peroxiredoxin | *bcp* | cellular response to oxidative stress | 0,574 |
| Q9HUG9 | Bifunctional protein HldE | *hldE* | biosynthetic process | 0,573 |
| Q9HV96 | Aminoglycoside phosphotransferase domain-containing protein | | | 0,571 |
| P42257 | Protein PilJ | *pilJ* | chemotaxis | 0,570 |
| Q9HTQ0 | D-amino acid dehydrogenase 1 | *dadA1* | catabolic process | 0,570 |
| Q9HXI4 | Nus factor SuhB | *suhB* | metabolic process | 0,570 |
| Q9I0K0 | NADH-quinone oxidoreductase subunit B | *nuoB* | aerobic respiration | 0,570 |
| Q9HXR8 | Probable FMN oxidoreductase | | oxidoreductase (mol) | 0,569 |
| G3XCV0 | Transcriptional regulator FleQ | *fleQ* | dna-templated transcription | 0,568 |
| Q51551 | Dihydroorotase-like protein | *pyrC'* | biosynthetic process | 0,567 |
| Q9HVZ9 | UDP-N-acetylmuramoylalanine--D-glutamate ligase | *murD* | cell wall organitation | 0,567 |
| Q9I5I4 | Probable enoyl-CoA hydratase/isomerase | | catabolism | 0,567 |
| Q9HUK1 | DNA topoisomerase 4 subunit A | *parC* | chromosome segregation | 0,566 |
| Q9I352 | Bacteriohemerythrin | | transport | 0,566 |
| Q9I1N4 | PslE | *pslE* | biofilm formation | 0,564 |
| Q9I5E2 | 2-methylisocitrate lyase | *prpB* | catabolic process | 0,564 |
| Q9HWG0 | UvrABC system protein A | *uvrA* | dna damage | 0,564 |

| | | | | |
|---|---|---|---|---|
| Q9I137 | Glycine dehydrogenase (decarboxylating) 1 | *gcvP1* | oxidoreductase (mol) | 0,562 |
| Q9HZK6 | Na(+)-translocating NADH-quinone reductase subunit A | *nqrA* | ion transport | 0,561 |
| Q9I669 | HotDog ACOT-type domain-containing protein | | metabolic process | 0,560 |
| Q9HXY7 | 3-hydroxyacyl-[acyl-carrier-protein] dehydratase FabZ | *fabZ* | lipid metabolism | 0,560 |
| Q9HZ66 | Phosphoserine aminotransferase | *serC* | amino-acid biosynthesis | 0,560 |
| Q9HV59 | Polyribonucleotide nucleotidyltransferase | *pnp* | catabolic process | 0,559 |
| Q9HXN2 | Phosphoribosylformylglycinamidine synthase | *purL* | purine biosynthesis | 0,557 |
| Q9HU31 | Amino acid ABC transporter | | | 0,556 |
| Q9HX91 | Uncharacterized protein PA3922 | | protein biosynthesis | 0,556 |
| Q9I4R0 | Lipoprotein | | | 0,555 |
| Q9I6H5 | D-3-phosphoglycerate dehydrogenase | *serA* | metabolism | 0,555 |
| Q9I6M5 | Glutarate-semialdehyde dehydrogenase | *davD* | catabolic process | 0,555 |
| Q9HXZ5 | Enolase | *eno* | glycolysis | 0,555 |
| Q9I502 | Proline--tRNA ligase | *proS* | protein biosynthesis | 0,554 |
| Q9HZA7 | Acetyl-coenzyme A carboxylase carboxyl transferase subunit beta | *accD* | fatty acid biosynthesis | 0,554 |
| Q9HUM3 | Protein HflC | *hflC* | proteolysis | 0,553 |
| Q9I0M6 | Serine--tRNA ligase | *serS* | protein biosynthesis | 0,553 |
| Q9HZE0 | NAD-specific glutamate dehydrogenase | *gdhB* | oxidoreductase (mol) | 0,553 |
| Q9I0K9 | Adenylosuccinate lyase | *purB* | biosynthetic process | 0,553 |
| P50599 | Tol-Pal system protein TolR | *tolR* | cell division/ protein transport | 0,552 |
| Q9HVT8 | Glutamyl-tRNA(Gln) amidotransferase subunit A | *gatA* | translation | 0,552 |
| Q9I2W9 | Phosphoenolpyruvate synthase | *ppsA* | gluconeogenesis | 0,552 |
| Q9I6M4 | 5-aminovalerate aminotransferase DavT | *davT* | catabolic process | 0,552 |

| | | | | |
|---|---|---|---|---|
| Q9HXH0 | Valine--tRNA ligase | *valS* | protein biosynthesis | 0,552 |
| G3XDB0 | Catabolite repression control protein | *crc* | quorum sensing | 0,551 |
| Q9HU22 | Glucose-1-phosphate thymidylyltransferase | *rmlA* | biosynthetic process | 0,550 |
| P08280 | Protein RecA | *recA* | dna binding | 0,549 |
| P23189 | Glutathione reductase | *gor* | cell redox homeostasis | 0,549 |
| P29365 | Homoserine dehydrogenase | *hom* | biosynthetic process | 0,549 |
| Q9I672 | CoA transferase | | transferase (mol) | 0,549 |
| Q9I099 | Threonine--tRNA ligase | *thrS* | protein biosynthesis | 0,549 |
| P0C2B2 | Thiol:disulfide interchange protein DsbA | *dsbA* | oxidoreductase activity | 0,548 |
| P55222 | cAMP-activated global transcriptional regulator Vfr | *vfr* | dna binding | 0,545 |
| Q9I5E4 | aconitate hydratase | | metabolic process | 0,545 |
| Q9I1N5 | PslD | *pslD* | biofilm formation | 0,544 |
| Q9HWR2 | Aminoglycoside 3'-phosphotransferase | *aph* | antibiotic resistance | 0,544 |
| Q9I0J9 | NADH-quinone oxidoreductase subunit C/D | *nuoC* | aerobic respiration | 0,544 |
| P55218 | O-succinylhomoserine sulfhydrylase | *metZ* | biosynthetic process | 0,543 |
| Q9I2P8 | Fe/S biogenesis protein NfuA | *nfuA* | iron binding (mol) | 0,543 |
| Q9I6B4 | Thiazole synthase | *thiG* | biosynthesis | 0,543 |
| Q9I0J7 | NADH-quinone oxidoreductase subunit F | *nuoF* | aerobic respiration | 0,542 |
| Q9HVX6 | Tryptophan--tRNA ligase | *trpS* | tryptophanyl-trna aminoacylation | 0,541 |
| Q9I296 | Isovaleryl-CoA dehydrogenase, mitochondrial | *liuA* | catabolic process | 0,541 |
| Q9I649 | Protein phosphatase 2C domain-containing protein | | | 0,540 |
| O50174 | N-succinylglutamate 5-semialdehyde dehydrogenase | *astD* | catabolic process | 0,539 |
| Q14T74 | proton-translocating NAD(P)(+) transhydrogenase | *pntAA* | nadph regeneration | 0,539 |

| | | | | |
|---|---|---|---|---|
| Q9I6G0 | L-threonine dehydratase | *ilvA1* | amino-acid biosynthesis | 0,539 |
| Q9HTF1 | Low specificity L-threonine aldolase | *ltaE* | biosynthetic process | 0,538 |
| Q9HVA0 | Acetolactate synthase | *ilvI* | biosynthetic process | 0,538 |
| Q9I140 | aminomethyltransferase | *gcvT2* | catabolic process | 0,538 |
| Q9I2T8 | Periplasmic chaperone PpiD | *ppiD* | isomerase (mol) | 0,538 |
| Q9I2U7 | Cysteine--tRNA ligase | *cysS* | protein biosynthesis | 0,537 |
| Q9HYT1 | Solute-binding protein family 3/N-terminal domain-containing protein | | transmembrane transport | 0,537 |
| G3XD24 | Methyl-accepting chemotaxis protein PctA | *pctA* | chemotaxis | 0,534 |
| Q9HWC4 | Transcription termination/antitermination protein NusG | *nusG* | dna-templated transcription termination | 0,534 |
| Q9I4W0 | Phosphoribosylaminoimidazole-succinocarboxamide synthase | *purC* | purine biosynthesis | 0,534 |
| Q9I6H6 | DUF4399 domain-containing protein | | unknown | 0,534 |
| Q9HYD5 | NAD-dependent malic enzyme | *maeA* | metabolic process | 0,534 |
| P43903 | Quinone oxidoreductase | *qor* | zinc ion binding | 0,533 |
| Q9I3L9 | Thiosulfate-binding protein | *cysP* | transport | 0,533 |
| Q9I526 | Cysteine synthase B | *cysM* | stringent response | 0,533 |
| Q9HWE2 | Large ribosomal subunit protein uL16 | *rplP* | translation | 0,532 |
| Q9HWF5 | Protein translocase subunit SecY | *secY* | protein transport | 0,532 |
| Q9HXY1 | Methionine aminopeptidase | *map* | proteolysis | 0,531 |
| Q9HTT8 | HemY N-terminal domain-containing protein | | metabolic process | 0,530 |
| Q9HX20 | Gamma-glutamyl phosphate reductase | *proA* | amino-acid biosynthesis | 0,530 |
| Q9I2S4 | Probable enoyl-CoA hydratase/isomerase | | fatty acid metabolism | 0,528 |
| Q9I2U1 | ATP-dependent Clp protease proteolytic subunit 1 | *clpP1* | cellular response to antibiotic | 0,528 |
| Q9I1M2 | 2-oxoisovalerate dehydrogenase subunit alpha | *bkdA1* | catabolic process | 0,526 |

| | | | | |
|---|---|---|---|---|
| Q9HX44 | Probable acyl-CoA dehydrogenase | | oxidoreductase (mol) | 0,526 |
| Q9I083 | Probable outer membrane protein | | | 0,526 |
| Q9I0A4 | Phenylalanine--tRNA ligase beta subunit | *pheT* | protein biosynthesis | 0,526 |
| Q9HVZ7 | UDP-N-acetylmuramoyl-tripeptide--D-alanyl-D-alanine ligase | *murF* | cell wall organitation | 0,525 |
| Q9I1Z6 | Alcohol dehydrogenase (Zn-dependent) | | oxidoreductase (mol) | 0,525 |
| Q9HXG8 | ABC transporter substrate-binding protein | | | 0,525 |
| Q9I0A3 | Phenylalanine--tRNA ligase alpha subunit | *pheS* | protein biosynthesis | 0,524 |
| Q9HVU9 | PmbA protein | *pmbA* | proteolysis | 0,523 |
| Q9HW15 | Probable short-chain dehydrogenase | *speA* | fatty acid elongation | 0,523 |
| Q9HW02 | UDP-N-acetylmuramate--L-alanine ligase | *murC* | cell wall organitation | 0,522 |
| Q9LCT3 | Protein translocase subunit SecA | *secA* | protein transport | 0,521 |
| Q9HTJ1 | NAD/NADP-dependent betaine aldehyde dehydrogenase | *betB* | aldehyde dehydrogenase | 0,520 |
| Q9HU15 | Beta-ketoacyl-[acyl-carrier-protein] synthase FabY | *fabY* | biosynthetic process | 0,520 |
| Q9XCL6 | Glutamate--tRNA ligase | *gltX* | protein biosynthesis | 0,520 |
| Q9I067 | FAD dependent oxidoreductase domain-containing protein | | oxidoreductase (mol) | 0,519 |
| Q9I0J4 | NADH-quinone oxidoreductase subunit I | *nuoI* | aerobic respiration | 0,519 |
| P72173 | Aspartate aminotransferase | *aspC* | biosynthetic process | 0,518 |
| Q9HXK5 | 2-isopropylmalate synthase | *leuA* | amino-acid biosynthesis | 0,518 |
| Q9I5A4 | Acetate kinase | *ackA* | metabolic process | 0,517 |
| Q9HUV9 | Bifunctional purine biosynthesis protein PurH | *purH* | biosynthetic process | 0,516 |
| Q9I2U0 | ATP-dependent Clp protease ATP-binding subunit ClpX | *clpX* | catabolic process | 0,516 |
| Q9I5U3 | Chaperone SurA | *surA* | protein folding | 0,515 |
| Q9HUE4 | Homocysteine synthase | *metY* | biosynthetic process | 0,514 |

| | | | | |
|---|---|---|---|---|
| Q9HZP5 | Electron transfer flavoprotein-ubiquinone oxidoreductase | | electron transport | 0,514 |
| P38103 | 4-hydroxy-tetrahydrodipicolinate reductase | *dapB* | biosynthetic process | 0,513 |
| Q9HWE5 | Large ribosomal subunit protein uL14 | *rplN* | translation | 0,513 |
| Q9HY11 | AMP nucleosidase | | | 0,513 |
| Q9I612 | Probable acyl-CoA dehydrogenase | | oxidoreductase (mol) | 0,512 |
| Q9I6J1 | Putrescine-binding periplasmic protein SpuD | *spuD* | transport | 0,512 |
| Q9ZFK4 | 2-dehydro-3-deoxyphosphooctonate aldolase | *kdsA* | biosynthesis | 0,512 |
| G3XD76 | Tetrahydropicolinate succinylase | *dapD* | biosynthetic process | 0,511 |
| P24474 | Nitrite reductase | *nirS* | electron transport | 0,510 |
| Q9I1N7 | mannose-1-phosphate guanylyltransferase | *pslB* | biofilm formation | 0,510 |
| Q9I6A7 | DUF4426 domain-containing protein | | unknown | 0,510 |
| Q9I7B7 | Glycine--tRNA ligase alpha subunit | *glyQ* | protein biosynthesis | 0,510 |
| Q9I0J8 | NADH-quinone oxidoreductase subunit E | *nuoE* | aerobic respiration | 0,509 |
| Q9I777 | DUF1993 domain-containing protein | | unknown | 0,508 |
| Q9HY84 | Argininosuccinate synthase | *argG* | amino-acid biosynthesis | 0,508 |
| O69077 | Aspartokinase | *lysC* | biosynthetic process | 0,507 |
| P72158 | N5-carboxyaminoimidazole ribonucleotide synthase | *purK* | biosynthetic process | 0,507 |
| Q9HUV8 | Phosphoribosylamine--glycine ligase | *purD* | biosynthetic process | 0,507 |
| P50987 | Argininosuccinate lyase | *argH* | biosynthetic process | 0,506 |
| Q51344 | Aspartate-semialdehyde dehydrogenase | *asd* | biosynthetic process | 0,506 |
| Q9HU38 | AsmA domain-containing protein | | regulation of protein targeting to membrane | 0,506 |
| Q9HZJ2 | Fatty acid oxidation complex subunit alpha | *fadB* | fatty acid metabolism | 0,506 |
| Q9HU43 | imidazole-4-carboxamide isomerase | *hisA* | biosynthetic process | 0,505 |

| | | | | |
|---|---|---|---|---|
| Q9HUA5 | Glucans biosynthesis protein G | *opgG* | metabolic process | 0,505 |
| Q9HTQ2 | Alanine racemase, catabolic | *dadX* | peptidoglycan biosynthetic process | 0,504 |
| Q9I2Y7 | Phospho-2-dehydro-3-deoxyheptonate aldolase | | amino-acid biosynthesis | 0,504 |
| Q51422 | Aspartate--tRNA(Asp/Asn) ligase | *aspS* | atp binding | 0,503 |
| Q9I003 | ATP-dependent RNA helicase DeaD | *deaD* | catabolic process | 0,503 |
| Q9HXJ5 | Histidine--tRNA ligase | *hisS* | protein biosynthesis | 0,502 |
| Q9HYV7 | Beta-ketodecanoyl-[acyl-carrier-protein] synthase | | biosynthetic process | 0,502 |
| Q9HUK5 | Phosphoserine phosphatase | | biosynthetic process | 0,501 |
| G3XD72 | Probable purine-binding chemotaxis protein | | chemotaxis | 0,500 |
| Q9HVF1 | Probable malate:quinone oxidoreductase 2 | *mqo2* | oxidoreductase activity | 0,499 |
| Q51342 | Amidophosphoribosyltransferase | *purF* | biosynthetic process | 0,498 |
| Q9I3I5 | Cell division protein ZipA | *zipA* | cell cycle-cell division | 0,498 |
| Q9HY12 | MBL fold metallo-hydrolase | | snrna processing | 0,498 |
| Q9HZC5 | Aminopeptidase N | *pepN* | proteolysis | 0,498 |
| Q9I271 | Cysteine protease | | proteolysis | 0,496 |
| Q9I2Q7 | Sulfite reductase | *cysI* | iron binding (mol) | 0,496 |
| Q9HXI0 | Sec translocon accessory complex subunit YajC | | protein transport | 0,496 |
| O52761 | Large ribosomal subunit protein bL17 | *rplQ* | translation | 0,495 |
| Q9I6J0 | Spermidine-binding periplasmic protein SpuE | *spuE* | transport-virulence | 0,495 |
| P77915 | oxidase | *hemN* | biosynthetic process | 0,494 |
| Q9HW72 | Pyruvate kinase | *pykA* | glycolitic process | 0,494 |
| P47203 | Cell division protein FtsA | *ftsA* | cell division | 0,493 |
| Q9I2U8 | Glutamine--tRNA ligase | *glnS* | protein biosynthesis | 0,493 |

| | | | | |
|---|---|---|---|---|
| Q9HYZ6 | Septum site-determining protein MinD | *minD* | cell division | 0,493 |
| P11436 | Aliphatic amidase | *amiE* | catabolic process | 0,492 |
| G3XD20 | Probable periplasmic serine endoprotease DegP-like | *mucD* | protein quality control | 0,491 |
| Q9I4S5 | Pyridoxine/pyridoxamine 5'-phosphate oxidase | *pdxH* | pyridoxine biosynthesis | 0,490 |
| P48247 | Glutamate-1-semialdehyde 2,1-aminomutase | *hemL* | biosynthetic process | 0,489 |
| Q9HT19 | ATP synthase gamma chain | *atpG* | atp synthesis | 0,489 |
| Q9HVU0 | Cell shape-determining protein MreB | *mreB* | regulation of cell shape | 0,489 |
| Q9I5Q9 | N-acetyl-gamma-glutamyl-phosphate reductase | *argC* | amino-acid biosynthesis | 0,489 |
| Q9I6J2 | Putrescine--pyruvate aminotransferase | *spuC* | biosynthetic process | 0,489 |
| Q9HXI8 | Cysteine desulfurase IscS | *iscS* | ion binding (mol) | 0,489 |
| Q59641 | Peptidyl-prolyl cis-trans isomerase A | *ppiA* | protein folding | 0,488 |
| P00106 | Cytochrome c4 | *cc4* | electron transport | 0,487 |
| Q9I7C4 | Beta sliding clamp | *dnaN* | dna replication | 0,486 |
| Q9HX33 | Leucine--tRNA ligase | *leuS* | protein biosynthesis | 0,486 |
| Q9HXE5 | ATP-dependent RNA helicase RhlB | *rhlB* | catabolic process | 0,486 |
| P38100 | Carbamoyl phosphate synthase large chain | *carB* | biosynthetic process | 0,485 |
| Q9HU83 | Urocanate hydratase | *hutU* | catabolic process | 0,485 |
| Q9I2T9 | Lon protease | *lon* | cellular response to antibiotic | 0,485 |
| Q9HZZ0 | L,D-TPase catalytic domain-containing protein | | cell shape | 0,485 |
| Q9HUM6 | Adenylosuccinate synthetase | *purA* | biosynthetic process | 0,484 |
| Q9I0T2 | Probable acyl-CoA dehydrogenase | | catabolic process | 0,483 |
| Q9X6R0 | Polyamine aminopropyltransferase 1 | *speE1* | biosynthesis | 0,482 |
| Q9HX46 | AMP nucleosidase | *amn* | metabolic process | 0,482 |

| | | | | |
|---|---|---|---|---|
| Q9HYU4 | Long-chain-fatty-acid--CoA ligase | *fadD1* | catabolic process | 0,482 |
| Q9I558 | Acetyl-coenzyme A synthetase 1 | *acsA1* | biosynthetic process | 0,481 |
| Q9HU53 | 2,3-bisphosphoglycerate-independent phosphoglycerate mutase | *gpmI* | metabolic process | 0,480 |
| Q9HZ63 | Ubiquinone biosynthesis O-methyltransferase | *ubiG* | biosynthetic process | 0,480 |
| Q9HVU7 | Metalloprotease TldD | | proteolysis | 0,479 |
| Q9I3G6 | Probable ferredoxin | | electron transport | 0,479 |
| Q9I5I5 | 3-hydroxyisobutyryl-CoA hydrolase | | catabolic process | 0,479 |
| P47205 | UDP-3-O-acyl-N-acetylglucosamine deacetylase | *lpxC* | biosynthetic process | 0,478 |
| Q9I6H4 | FAD-binding PCMH-type domain-containing protein | | biosynthetic process | 0,478 |
| Q9HVC2 | Ribosome-binding ATPase Ych | *ychF* | ribosomal large subunit binding | 0,477 |
| P50601 | Tol-Pal system protein TolB | *tolB* | cell division/ protein import | 0,476 |
| Q9HU41 | Imidazoleglycerol-phosphate dehydratase | *hisB* | biosynthetic process | 0,476 |
| Q9I2R2 | Probable oxidoreductase | | oxidoreductase (mol) | 0,476 |
| Q9I576 | 4-hydroxyphenylpyruvate dioxygenase | *hpd* | catabolism | 0,476 |
| Q9HVL8 | GTPase Obg | *obg* | ribosome biogenesis | 0,475 |
| P57112 | Soluble pyridine nucleotide transhydrogenase | *sthA* | metabolic process | 0,474 |
| Q9I1F5 | Probable glyceraldehyde-3-phosphate dehydrogenase | | catabolic process | 0,474 |
| Q9I7C2 | DNA gyrase subunit B | *gyrB* | dna replication | 0,474 |
| Q9I0L2 | tRNA-specific 2-thiouridylase MnmA | *mnmA* | trna processing | 0,474 |
| G3XD74 | serine-type D-Ala-D-Ala carboxypeptidase | *dacC* | proteolysis | 0,473 |
| Q9HUM7 | Ribonuclease R | *rnr* | catabolic process | 0,473 |
| Q9I3D8 | Uncharacterized protein PA1579 | | lipid binding (mol) | 0,472 |
| Q9HT06 | Membrane protein insertase YidC | *yidC* | protein transport | 0,471 |

| | | | | |
|---|---|---|---|---|
| Q9HTX5 | Aminomethyltransferase | *gcvT* | transaminase activity | 0,471 |
| P14165 | Citrate synthase | *gltA* | tricarboxylic acid cycle | 0,470 |
| Q9I553 | Alanine--tRNA ligase | *alaS* | protein biosynthesis | 0,470 |
| Q9I5Y8 | Transketolase | *tktA* | transferase (mol) | 0,469 |
| Q51485 | Porin B | *oprB* | porin activity | 0,468 |
| Q9HU50 | Probable carboxyl-terminal protease | | protein secretion | 0,468 |
| Q9HVW8 | ATP phosphoribosyltransferase | *hisG* | biosynthetic process | 0,468 |
| Q9I1M0 | Lipoamide acyltransferase component of branched-chain alpha-keto acid dehydrogenase complex | *bkdB* | glycolysis | 0,466 |
| Q9I344 | Chorismate synthase | *aroC* | amino-acid biosynthesis | 0,466 |
| Q9I5U2 | LPS-assembly protein LptD | *lptD* | biosynthetic process | 0,466 |
| Q9HZP8 | Enoyl-[acyl-carrier-protein] reductase [NADH] | *fabV* | fatty acid biosynthesis | 0,466 |
| P26276 | Phosphomannomutase | *algC* | biosynthetic process | 0,465 |
| Q9HUQ6 | Probable short-chain dehydrogenase | | oxidoreductase activity | 0,464 |
| Q51390 | Glycerol kinase 2 | *glpK2* | biosynthetic process | 0,462 |
| Q9HT76 | Vitamin B12-dependent ribonucleotide reductase | *nrdJa* | biosynthetic process | 0,462 |
| Q9HW04 | Arginine biosynthesis bifunctional protein ArgJ | *argJ* | biosynthetic process | 0,462 |
| Q9I456 | Probable outer membrane protein | | | 0,461 |
| Q9I298 | 3-methylglutaconyl-CoA hydratase | *liuC* | catabolic process | 0,460 |
| Q9I6J9 | Agmatine deiminase | *aguA* | biosynthesis | 0,460 |
| P0DP44 | Polyphosphate kinase | *ppk* | biosynthetic process | 0,459 |
| Q9I5F9 | Lon protease | *lon* | cellular response to antibiotic | 0,459 |
| Q9HUF1 | Peptide methionine sulfoxide reductase MsrA | *msrA* | cellular response | 0,458 |
| Q9HZ15 | Probable transcriptional regulator | | transcription regulation | 0,458 |

| | | | | |
|---|---|---|---|---|
| Q9HVT2 | Alpha-2-macroglobulin homolog | | endopeptidase inhibitor activity | 0,457 |
| Q9HX32 | LPS-assembly lipoprotein LptE | *lptE* | transport | 0,456 |
| P72170 | Dihydroorotase | *pyrC* | biosynthetic process | 0,455 |
| Q9HY01 | S-(hydroxymethyl)glutathione dehydrogenase | *adhC* | catabolic process | 0,455 |
| O68822 | Cytosol aminopeptidase | *pepA* | rna metabolic process | 0,454 |
| Q9HUG0 | Probable carbamoyl transferase | | biosynthetic process | 0,452 |
| Q9HV50 | Phosphoglucosamine mutase | *glmM* | biosynthetic process | 0,452 |
| Q9HWD7 | Large ribosomal subunit protein uL23 | *rplW* | translation | 0,452 |
| Q9I513 | Phosphoribosylformylglycinamidine cyclo-ligase | *purM* | purine biosynthesis | 0,452 |
| Q9I7A7 | LysM domain-containing protein | | | 0,452 |
| Q9HZL1 | Na(+)-translocating NADH-quinone reductase subunit F | *nqrF* | ion transport | 0,451 |
| Q9I0J6 | NADH-quinone oxidoreductase subunit G | *nuoG* | aerobic respiration | 0,451 |
| Q9HU18 | C4-dicarboxylate-binding periplasmic protein DctP | *dctP* | transmembrane transport | 0,450 |
| P52477 | Multidrug resistance protein MexA | *mexA* | response to antibiotic/ transmembrane transport | 0,449 |
| Q9HVJ1 | Energy-dependent translational throttle protein EttA | *ettA* | translation | 0,449 |
| O33877 | 3-hydroxydecanoyl-[acyl-carrier-protein] dehydratase | *fabA* | fatty acid biosynthetic process | 0,448 |
| Q9HTV1 | Transcription termination factor Rho | *rho* | rna binding | 0,448 |
| Q9I4L6 | Outer-membrane lipoprotein YfiB | *yfiB* | | 0,448 |
| Q9I2X2 | Inactive transglutaminase fused to 7 transmembrane helices | | | 0,446 |
| O50177 | Succinylglutamate desuccinylase | *astE* | catabolic process | 0,445 |
| P28810 | Methylmalonate-semialdehyde dehydrogenase | *mmsA* | catabolic process | 0,445 |
| O86428 | Branched-chain-amino-acid aminotransferase | *ilvE* | biosynthetic process | 0,444 |
| Q9I028 | Acyl-coenzyme A dehydrogenase | | fatty acid biosynthesis | 0,443 |

| | | | | |
|---|---|---|---|---|
| P26275 | Positive alginate biosynthesis regulatory | *algR* | dna binding | 0,442 |
| Q9HVY9 | BON domain-containing protein | | response to osmotic stress | 0,442 |
| Q9I2A2 | fumarylacetoacetase | *fahA* | catabolism | 0,442 |
| Q9I3F6 | Methyl-accepting chemotaxis protein Aer | *aer* | chemotaxis | 0,442 |
| Q9I747 | Protein hcp1 | *hcp1* | protein binding (mol) | 0,441 |
| Q9HV48 | ATP-dependent zinc metalloprotease FtsH | *ftsH* | cell division | 0,440 |
| Q9HW49 | UspA domain-containing protein | | | 0,440 |
| Q9I6E0 | Dihydroxy-acid dehydratase | *ilvD* | amino-acid biosynthesis | 0,440 |
| G3XCU6 | Transcriptional regulator CysB | *cysB* | regulation of dna-templated transcription | 0,439 |
| P48372 | DNA gyrase subunit A | *gyrA* | dna binding | 0,438 |
| Q9HXI1 | Protein translocase subunit SecD | *secD* | protein transport | 0,436 |
| Q9HU67 | Large ribosomal subunit assembly factor BipA | *typA* | biofilm formation, response to antibiotic | 0,434 |
| Q9I407 | Glutaminase-asparaginase | *ansB* | catabolic process | 0,434 |
| Q9I1F8 | Probable oxidoreductase | | oxidoreductase (mol) | 0,431 |
| P52002 | Multidrug resistance protein MexB | *mexB* | response to antibiotic/ transmembrane transport | 0,430 |
| Q9HXZ2 | Acetyl-coenzyme A carboxylase carboxyl transferase subunit alpha | *accA* | lipid metabolism | 0,430 |
| Q9HZF8 | Dihydroorotate dehydrogenase (quinone) | *pyrD* | pyrimidine biosynthesis | 0,429 |
| Q9HYI7 | Leucine dehydrogenase | *ldh* | metabolic process | 0,428 |
| Q9HXZ4 | CTP synthase | *pyrG* | biosynthesis | 0,427 |
| Q9HTV8 | Membrane fusion protein biotin-lipoyl | | no function | 0,426 |
| Q9I658 | DUF2780 domain-containing protein | | unknown | 0,426 |
| Q9HW87 | Formyltetrahydrofolate deformylase | *purU1* | biosynthetic process | 0,425 |

| | | | | |
|---|---|---|---|---|
| Q9HVB9 | Carbonic anhydrase | | cellular response | 0,424 |
| O50175 | N-succinylarginine dihydrolase | *astB* | catabolic process | 0,423 |
| P52111 | Glycerol-3-phosphate dehydrogenase | *glpD* | metabolic process | 0,422 |
| Q9HXP9 | Small ribosomal subunit protein bS16 | *rpsP* | translation | 0,422 |
| Q9I0I4 | Methyl-accepting chemotaxis protein TlpQ | *tlpQ* | chemotaxis | 0,420 |
| G3XCT6 | Chemotaxis protein CheA | | chemotaxis | 0,419 |
| Q9HUW8 | DUF2333 family protein | | | 0,419 |
| O33407 | Esterase EstA | *estA* | cell motility | 0,416 |
| Q9HZJ3 | 3-ketoacyl-CoA thiolase | *fadA* | fatty acid metabolism | 0,416 |
| Q9HTY8 | S1 motif domain-containing protein | | translation | 0,415 |
| Q9HUJ8 | DNA topoisomerase 4 subunit B | *parE* | chromosome segregation | 0,414 |
| P28811 | 3-hydroxyisobutyrate dehydrogenase | *mmsB* | catabolic process | 0,412 |
| P72157 | N5-carboxyaminoimidazole ribonucleotide mutase | *purE* | biosynthetic process | 0,411 |
| Q9HXJ7 | Outer membrane protein assembly factor BamB | *bamB* | protein transport | 0,410 |
| Q9I1L9 | Dihydrolipoyl dehydrogenase | *lpdV* | oxidoreductase (mol) | 0,405 |
| Q9HYL2 | Nitrous-oxide reductase | *nosZ* | oxidoreductase (mol) | 0,405 |
| Q9I3G5 | Cbb3-type cytochrome c oxidase subunit | *ccoP1* | electron transport | 0,404 |
| Q9HWS8 | Probable dehydrogenase | | biosynthetic process | 0,403 |
| Q9I1I5 | Glucose dehydrogenase | *gcd* | oxidoreductase (mol) | 0,401 |
| Q9HXB0 | Peptide chain release factor 3 | *prfC* | protein biosynthesis | 0,400 |
| Q9I0T0 | Probable short-chain dehydrogenase | | metabolic processes | 0,393 |
| Q9I2A9 | 3-oxoadipate CoA-transferase | *dhcB* | catabolic process | 0,391 |
| Q9I5G8 | Elongation factor 4 | *lepA* | protein biosynthesis | 0,388 |

| | | | | |
|---|---|---|---|---|
| Q9I4T3 | Probable outer membrane protein | | monoatomic ion transport | 0,387 |
| Q9HVW7 | UDP-N-acetylglucosamine 1-carboxyvinyltransferase | *murA* | cell wall organization | 0,379 |
| Q9HY79 | Bacterioferritin | *bfrB* | ion transport | 0,378 |
| Q9HXI2 | Protein translocase subunit SecF | *secF* | protein transport | 0,377 |
| O82852 | Uridylate kinase | *pyrH* | biosynthetic process | 0,370 |
| Q9HVW0 | Arabinose 5-phosphate isomerase KdsD | *kdsD* | biosynthetic process | 0,357 |
| Q9HWF8 | Small ribosomal subunit protein uS11 | *rpsK* | translation | 0,353 |
| Q9HXY4 | Outer membrane protein assembly factor BamA | *opr86* | protein insertion into membrane | 0,353 |
| Q9HX05;Q9I2C4 | Probable aldehyde dehydrogenase | | oxidoreductase (mol) | 0,345 |
| G3XCX1 | nitrate reductase (quinone) | *narG* | nitrate metabolic process | 0,343 |
| Q9I689 | ATP-dependent RNA helicase RhlE | *rhlE* | ribosome biogenesis | 0,330 |
| Q9HWE8 | Small ribosomal subunit protein uS14 | *rpsN* | translation | 0,321 |
| P72174 | UvrABC system protein B | *uvrB* | dna binding | 0,275 |
| Q9X4G0 | Homogentisate 1,2-dioxygenase | *hmgA* | catabolism | 0,267 |
| Q9I702 | Malonate-semialdehyde dehydrogenase | *bauC* | biosynthetic process | 0,266 |
| Q9HZ67 | Bifunctional chorismate mutase/prephenate dehydratase | *pheA* | amino-acid biosynthesis | 0,266 |
| Q9I406 | Glutathione hydrolase proenzyme | *ggt* | glutathione biosynthesis | 0,252 |
| Q9HVM9 | Type 4 fimbrial biogenesis protein PilX | *pilX* | pilus assembly | 0,250 |
| Q9HTL5 | NAD-dependent epimerase/dehydratase domain-containing protein | | aldehyde dehydrogenase | 0,237 |
| Q9HYZ3 | Probable M18 family aminopeptidase 2 | *apeB* | proteolysis | 0,230 |
| Q51480 | Protein NirF | *nirF* | biosynthesis of heme d1 of nitrite reductase | 0,228 |
| Q9HVK8 | endopeptidase La | | proteolysis | 0,227 |
| Q9I524 | GTP pyrophosphokinase | *relA* | stringent response | 0,215 |

| | | | | |
|---|---|---|---|---|
| G3XDA2 | 3-oxoacyl-[acyl-carrier-protein] synthase 2 | *fabF1* | fatty acid biosynthetic process | 0,191 |
| Q9HT16 | ATP synthase subunit b | *atpF* | atp synthesis | 0,000 |
| Q9HT22 | Bifunctional protein GlmU | *glmU* | cell wall organization | 0,000 |
| Q9HT80 | DNA polymerase I | *polA* | dna binding | 0,000 |
| Q9HTN7 | Probable binding protein component of ABC dipeptide transporter | | dipeptide transport | 0,000 |
| Q9HUA6 | Glucans biosynthesis glucosyltransferase H | *opgH* | metabolic process | 0,000 |
| Q9HV69 | Pantothenate synthetase | *panC* | biosynthetic process | 0,000 |
| Q9HVV9 | 3-deoxy-D-manno-octulosonate 8-phosphate phosphatase KdsC | | biosynthetic process | 0,000 |
| Q9HVW9 | Histidinol dehydrogenase | *hisD* | biosynthetic process | 0,000 |
| Q9I297 | Methylcrotonyl-CoA carboxylase, beta-subunit | *liuB* | catabolic process | 0,000 |
| Q9I2B0 | 3-oxoadipate CoA-transferase | *dhcA* | catabolic process | 0,000 |
| Q9I2U6 | Bifunctional protein FolD | *folD* | amino-acid biosynthesis | 0,000 |
| Q9I3N3 | Cytochrome c-type biogenesis protein CcmE | *ccmE* | cytochrome c-type biogenesis | 0,000 |
| Q9I5H3 | Probable two-component response regulator | | transcription | 0,000 |
| Q9HWZ6 | Inorganic pyrophosphatase | *ppa* | ion binding (mol) | 0,000 |
| Q9HXJ8 | GTPase Der | *der* | ribosome biogenesis | 0,000 |
| Q9HZA3 | 3-isopropylmalate dehydratase large subunit | *leuC* | amino-acid biosynthesis | 0,000 |
| Q9I0M2 | Thioredoxin reductase | *trxB1* | oxidoreductase (mol) | 0,000 |
